# The HUSH complex controls brain architecture and protocadherin fidelity

**DOI:** 10.1101/2021.11.29.466909

**Authors:** Astrid Hagelkruys, Marion Horrer, Jasmin Taubenschmid-Stowers, Anoop Kavirayani, Maria Novatchkova, Michael Orthofer, Tsung-Pin Pai, Domagoj Cikes, Sergei Zhuk, Meritxell Balmaña, Christopher Esk, Rubina Koglgruber, Shane J.F. Cronin, Ulrich Elling, Jürgen A. Knoblich, Josef M. Penninger

## Abstract

Fine-tuning of neural connectivity is important for cerebral functions and brain evolution. Protocadherins provide barcodes for neuronal identity as well as synapse formation and expansion of protocadherin cluster genes has been linked to advanced cognitive functions. The tightly controlled stochastic and combinatorial expression of the different protocadherin isoforms in individual neurons provides the molecular basis for neuronal diversity, neuronal network complexity and function of the vertebrate brain. How protocadherins are epigenetically controlled has not yet been fully elucidated. Here we show that the HUSH (human silencing hub) complex containing H3K9me3 binding protein M-phase phosphoprotein 8 (MPP8) and Microrchidia CW-type zinc finger protein 2 (MORC2), critically controls the fidelity of protocadherin expression. MPP8 and MORC2A are highly expressed in the murine brain and exclusively found in neurons. Genetic inactivation of *Mphosph8* (coding for MPP8) or *Morc2a* in the nervous system of mice leads to increased brain size, altered brain architecture, and behavioral changes. Mechanistically, MPP8 and MORC2A precisely and selectively suppress the repetitive-like protocadherin gene cluster on mouse chromosome 18 in a H3K9me3-dependent manner, thereby affecting synapse formation. Moreover, we demonstrate that individual MPHOSPH8- or MORC2-deficient neurons in human cerebral organoids express increased numbers of clustered protocadherin isoforms. Our data identify the HUSH complex, previously linked to silencing of repetitive transposable elements, as a key epigenetic regulator of protocadherin expression in the nervous system and thereby brain development and neuronal individuality in mice and humans.

## Introduction

Histone methylation plays a crucial role in heterochromatin formation and is controlled by the opposing actions of “writers” (histone methyltransferases) and “erasers” (histone demethylases) ^1, 2^. “Reader” proteins bind to certain histone modifications and thereby influence fundamental biological processes such as transcription, replication or DNA repair. These epigenetic “writers”, “erasers” and “readers” need to be tightly regulated and disturbances in this highly balanced network of epigenetic control and chromatin-modifying enzymes can contribute to a variety of diseases including many types of cancers ^3^. Histone methylation has also been functionally linked to memory formation, neuroplasticity as well as cognitive capabilities and aberrant histone methylation has been associated with neurodevelopmental, neurodegenerative and behavioral disorders ^4–6^.

To identify factors involved in the repression of silenced elements, several genome-wide screens have been performed in human cell lines and identified the HUSH (human silencing hub) complex ^7–9^. This recently discovered repressor complex, which contains M-phase phosphoprotein 8 (MPP8) and recruits the histone methyltransferase SETDB1 as well as Microrchidia CW-type zinc finger protein 2 (MORC2), has been implicated in silencing of genes, repetitive elements, or transgenes in mammals ^7, 8^. Recently *MORC2* mutations have been reported in individuals with Charcot-Marie-Tooth (CMT) disease type 2Z, a form of axonal neuropathy with progressive muscle weakness, atrophy and sensory impairment ^10–12^ and in a neurodevelopmental disorder with intellectual disability, growth retardation, microcephaly and variable craniofacial dysmorphism ^13^.

Here we report that murine MPP8 and MORC2A are highly expressed in the brain and exclusively found in neurons. Genetic inactivation of *Mphosph8* (coding for MPP8) or *Morc2a* in the nervous system of mice results in altered brain architecture, impaired motor functions, and reduced lifespan. Mechanistically, in both mouse brains and human cerebral organoids MPP8 and MORC2A suppress the repetitive-like protocadherin gene cluster, involved in neuronal individuality and brain evolution, in a H3K9me3-dependent manner. Our data uncover a novel role for the HUSH complex in the regulation of clustered protocadherins within the nervous system, thereby controlling brain development and neuronal fidelity in mice and humans.

## Results

### Murine MPP8 and MORC2A are required for embryonic development

To elucidate the *in vivo* roles of MPP8 and MORC2A, we established mouse models carrying conditional *Mphosph8* or *Morc2a* knock-out alleles. Exons 4 of *Mphosph8* as well as *Morc2a* were flanked by loxP sites (Extended Data Fig. 1a,b) and Cre recombinase-mediated removal of this exon leads to out-of-frame splicing, subjecting the resulting mRNA to nonsense-mediated decay. Southern blot analysis of embryonic stem (ES) cells (Extended Data Fig. 1c,d) identified correctly targeted clones, which were then used for blastocyst injections. Loss of *Mphosph8* or *Morc2a* in mouse ES cells resulted in no apparent changes in proliferation but were significantly impaired in their ability to form teratomas (Extended Data Fig. 1e,f). *Actin*-Cre-mediated full-body knock-out of *Mphosph8* or *Morc2a* in the mouse resulted in embryonic lethality (Extended Data Fig. 1g,h). In comparison to a recent study showing that *Mphosph8* knock-out mice display partial embryonic lethality with surviving *Mphosph8*^-/-^ mice developing normally apart from reduced body weight ^14^, in our study *Actin*-Cre-mediated loss of *Mphosph8* resulted in 100% embryonic lethality before embryonic day 11.5 (E11.5) and loss of *Morc2a* led to embryonic lethality around E13.5 (Extended Data Fig. 1i,j). These data show that MPP8 and MORC2A are critically required for teratoma formation and embryonic development.

### Deletion of *Mphosph8* or *Morc2a* in the nervous system results in elevated brain/body ratios and altered brain architecture

Based on high expression of both MPP8 and MORC2A proteins in the adult mouse brain (Extended Data Fig. 1k,l) and the known link between histone methylation, neuroplasticity, and cognitive abilities ^4, 5^, we focused on investigating their function in the nervous system. Therefore, we crossed homozygous floxed *Mphosph8* and *Morc2a* mice to *Nestin*-Cre expressing mice, effectively deleting *Mphosph8* or *Morc2a* in neuronal progenitors and post-mitotic neurons from E11.0 onwards. *Mphosph8*^fl/fl^ *Nestin*-Cre and *Morc2a*^fl/fl^ *Nestin*-Cre mice are hereafter referred to as *Mphosph8*^Δ/Δn^ and *Morc2a*^Δ/Δn^, respectively. Loss of *Mphosph8* and *Morc2a* was verified by quantitative RT-PCRs (Fig. 1a and Extended Data Fig. 2a) and immunoblots (Fig. 1b and Extended Data Fig. 2b) from whole brains of control *Mphosph8*^fl/fl^ or *Morc2a*^fl/fl^ and their respective *Mphosph8*^Δ/Δn^ or *Morc2a*^Δ/Δn^ littermates. *Nestin*-Cre-mediated *Mphosph8* or *Morc2a*-deficiency in the central nervous system resulted in reduced body sizes, body weights and survival (Fig. 1c,d and Extended Data Fig. 2c,d).

**Figure 1.**
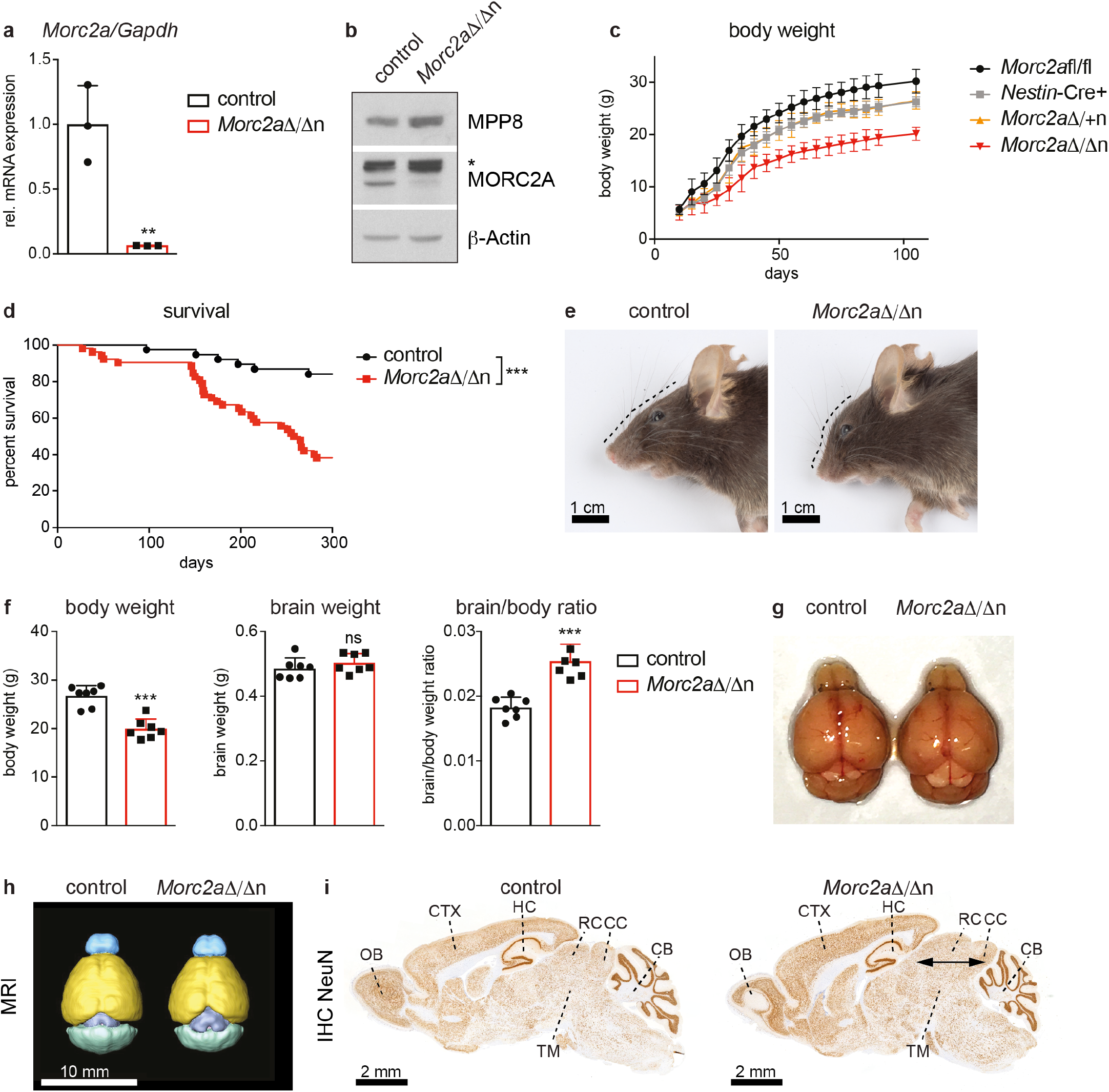
Altered brain architecture in *Morc2a* knock-out mice. (**a**) Relative mRNA expression of *Morc2a* in littermate control and homozygous *Morc2a*^Δ/Δn^ brains. Mean values +/- SD were normalized to the housekeeping gene *Gapdh* (n = 3). Data are shown for one of two independent experiments. (**b**) Immunoblot analysis of littermate control versus *Morc2a*^Δ/Δn^ whole brain extracts with antibodies against MPP8, MORC2A and β-Actin as loading control. The asterisk indicates an unspecific band. (**c**) Body weights of *Morc2a*^fl/fl^ (n = 13), *Nestin-*Cre+ (n = 8), heterozygous *Morc2a*^Δ/+n^ (n = 8) and *Morc2a*^Δ/Δn^ (n = 11) male littermates (+/- SD) during the first 3 months of age. Data from several litters were pooled. (**d**) Kaplan-Meier survival curve over the first 300 days of control versus *Morc2a*^Δ/Δn^ littermates. Data from several litters were pooled (n > 35) and survival curves were compared by the Log-rank (Mantel-Cox) test. (**e**) Representative macroscopic photographs of control (left) and *Morc2a*^Δ/Δn^ (right) mouse heads (scale bar = 1 cm). (**f**) Body weights, brain weights and brain/body ratios of control compared to *Morc2a*^Δ/Δn^ adult (at 3-4 months) littermates. Data from two independent experiments were pooled and are shown as mean values +/- SD (n = 7). (**g**) Representative macroscopic images of a control (left) and an *Morc2a*^Δ/Δn^ (right) littermate adult (at 3-4 months) brain. (**h**) Representative *in vivo* MRI brain scans of control (left) and *Morc2a*^Δ/Δn^ (right) adult (at 3-4 months) littermates (scale bar = 10 mm). In total 4 mice per genotype were analyzed. (**i**) NeuN immunohistochemistry (IHC) in adult (at 3-4 months) control (left) and *Morc2a*^Δ/Δn^ (right) brains (scale bar = 2 mm). In total at least 3 mice per genotype were analyzed by immunohistochemistry. OB = olfactory bulb, CTX = cerebral cortex, HC = hippocampus, RC = rostral colliculus, CC = caudal colliculus, TM = tegmentum, CB = cerebellum. Arrows indicate widening of collicular regions of the midbrain in *Morc2a*^Δ/Δn^ mice. For panels a and f each data point represents an individual mouse. *p* values were calculated using the Student’s *t* test. ** *p* < 0.01; *** *p* < 0.001; ns, not significant.

Morphologically, both *Mphosph8*^Δ/Δn^ and *Morc2a*^Δ/Δn^ mice featured varying degrees of cranial alterations including skulls that ranged from normal to moderately domed (Fig. 1e and Extended Data Fig. 2e). Intriguingly, nervous system-specific deletions of *Mphosph8* or *Morc2a* led to dramatically elevated brain/body ratios (Fig. 1f and Extended Data Fig. 2f). In both *Mphosph8*^Δ/Δn^ and *Morc2a*^Δ/Δn^ mice, the increase in brain mass corresponded to a partial macroencephaly featuring enlargement (dorsoventral and mediolateral expansion), predominantly of the collicular (tectal) and tegmental regions of the midbrain. This resulted in widening of the distance between the occipital lobes and the cerebellar vermis and imparted the appearance of exposed rostral and caudal colliculi. These changes were consistently observed by macroscopic examination (Fig. 1g and Extended Data Fig. 2g), Magnetic Resonance Imaging (MRI) (Fig. 1h and Extended Data Fig. 2h) as well as in histologic sections (Fig. 1i and Extended Data Fig. 2i). Quantitative immunohistochemistry revealed an increased density of RBFOX3+ (NeuN+) nuclei, especially in the colliculi (Extended Data Fig. 3a-d), indicative of increased numbers of neurons. Glial fibrillary acidic protein (GFAP) immunohistochemistry did not reveal any apparent differences in astrocyte numbers and distribution, suggesting that the observed morphologic expansion was attributable to increases in the neuronal rather than astrocytic components of the neuropil (data not shown). Of note, we found no histomorphological alterations in the olfactory bulb, cerebellar folia, the cerebral cortex or the hippocampal formation (Extended Data Fig. 3c-h). These data show that neuronal loss of murine MPP8 or MORC2A results in a morphological and neuronal expansion of defined brain areas.

### *Mphosph8* and *Morc2a* double mutant mice

To study if the loss of both MPHOSPH8 and MORC2A leads to more severe phenotypes, we generated *Mphosph8*^Δ/Δn^ *Morc2a*^Δ/Δn^ mice lacking both *Mphosph8* and *Morc2a* in the nervous system. Loss of *Mphosph8* and *Morc2a* was verified by quantitative RT-PCRs and immunoblots from whole brains of control (*Mphosph8*^fl/fl^ *Morc2a*^fl/fl^) and *Mphosph8*^Δ/Δn^ *Morc2a*^Δ/Δn^ littermates (Extended Data Fig. 4a,b). Upon *Nestin-*Cre-mediated double deletion of both *Mphosph8* and *Morc2a*, these mice showed elevated brain/body ratios and altered brain architecture including exposed colliculi, very similar to the changes observed in *Mphosph8*^Δ/Δn^ or *Morc2a*^Δ/Δn^ single mutant mice (Extended Data Fig. 4c-h). We observed a synergistic effect on mortality whereby survival rates of double mutant mice decreased after already 150 days to 50% and after 300 days to 20% (Extended Data Fig. 4i). These data indicate that, apart from enhanced lethality, double knock-out of *Mphosph8* and *Morc2a* leads to similar phenotypes in brain architecture as in single mutant mice, suggesting that these two molecules act in the same molecular complex.

### *Mphosph8* or *Morc2a* deficiency affects motor and memory functions

To evaluate the pathophysiological consequences of *Mphosph8* or *Morc2a* neuronal inactivation, we performed various behavioral tests analyzing locomotion, learning and memory in adult mice. Since *Mphosph8*^Δ/Δn^ and *Morc2a*^Δ/Δn^ mice showed the most notable neuroanatomic alterations in the midbrain, we paid special attention to motor control and coordination abilities. Both *Mphosph8* and *Morc2a* single knock-out mice exhibited decreased locomotor activity in PhenoMaster cages (Fig. 2a,b) and in the open field test (Fig. 2c,d). Moreover, the mutant mice exhibited reduced motor performance and impaired motor functions on the Rotarod test (Fig. 2e,f). To exclude *Nestin*-Cre specific effects, we performed similar experiments comparing *Nestin*-Cre- and *Nestin*-Cre+ mice and observed no significant differences in any of these assays (Extended Data Fig. 5a-c). *Mphosph8*^Δ/Δn^ and *Morc2a*^Δ/Δn^ mice also exhibited reduced neuromuscular function in the grip strength test (Fig. 2g,h), similar to the findings in human patients with *MORC2* mutations ^13^.

**Figure 2.**
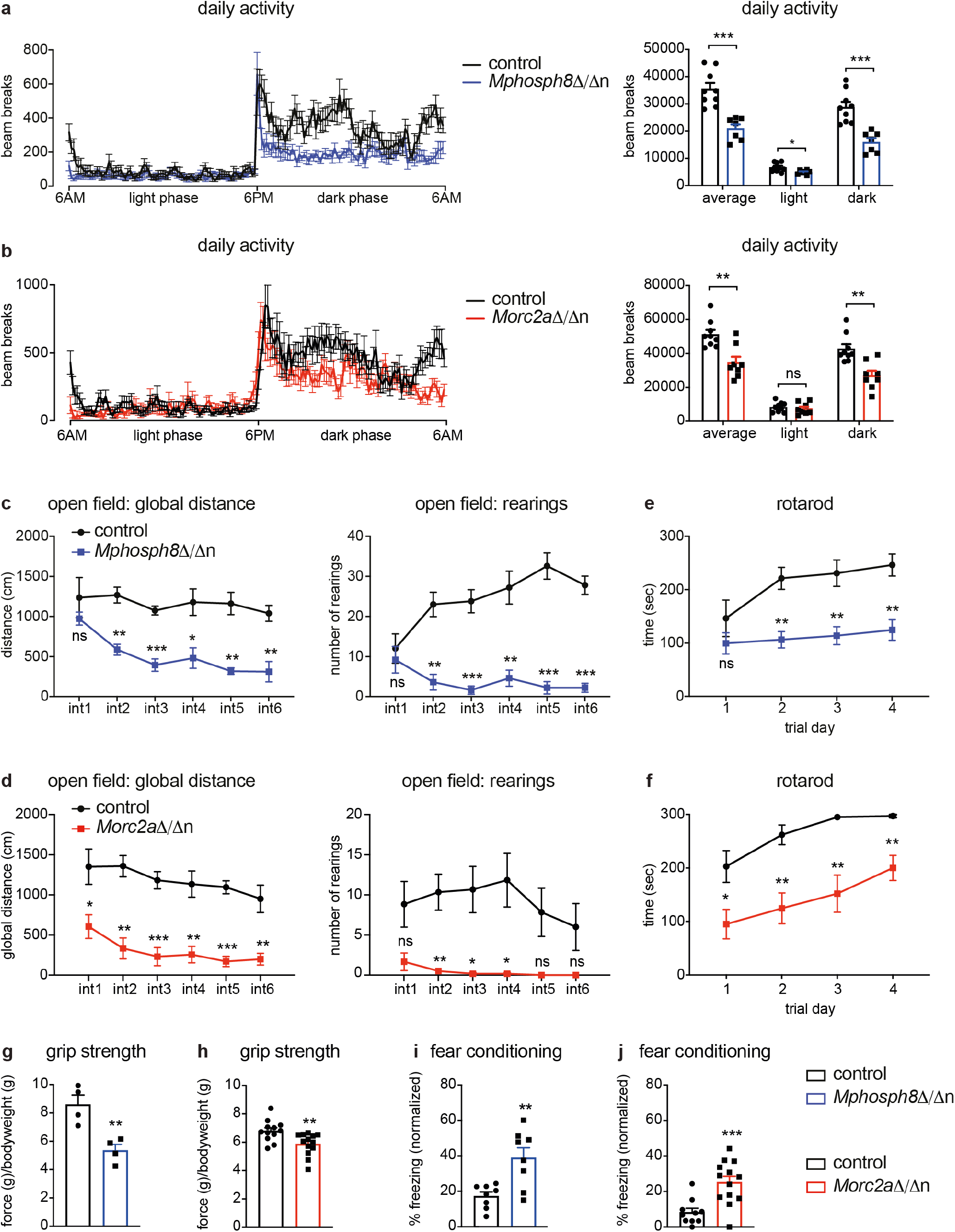
Decreased motor performance and enhanced fear memory upon loss of *Mphosph8* or *Morc2a*. (**a,b**) Daily activity in PhenoMaster cages measured by beam breaks of *Mphosph8*^Δ/Δn^ mice (**a,** n > 5) and *Morc2a*^Δ/Δn^ mice (**b**, n > 7) compared to their respective control littermates over 24 hours. Data from two independent experiments were pooled and shown as mean values +/- SEM. (**c,d**) Global distance and number of rearings of *Mphosph8*^Δ/Δn^ mice (**c**) and *Morc2a*^Δ/Δn^ mice (**d**) compared to their control littermates in the open field test. Data are shown as mean values +/- SEM (n > 5) and are representative of one of two independent experiments. (**e,f**) Latency to fall from the accelerating Rotarod of *Mphosph8*^Δ/Δn^ mice (**e**) and *Morc2a*^Δ/Δn^ mice (**f**) compared to control littermates. Data from two independent experiments were pooled and shown as mean values +/- SEM (n > 8). **(g,h)** 4-paw grip strength normalized to body weights of *Mphosph8*^Δ/Δn^ mice (**g**) and *Morc2a*^Δ/Δn^ mice (**h**) compared to control littermates. Data from two independent experiments were pooled and shown as mean values +/- SEM (n > 3). **(i,j)** Percent freezing in the fear-context box normalized to training without foot-shock of *Mphosph8*^Δ/Δn^ mice (**i**) and *Morc2a*^Δ/Δn^ mice (**j**) compared to control littermates. Data from two independent experiments were pooled and shown as mean values +/- SEM (n > 7). For the right panels in a,b and the panels in g-j each data point represents an individual mouse. *p* values were calculated using the Student’s *t* test. * *p* < 0.05; ** *p* < 0.01; *** *p* < 0.001; ns, not significant.

*Mphosph8*^Δ/Δn^ and *Morc2a*^Δ/Δn^ mice did not show any changes in the spontaneous alternation performance in the Y-maze as well as anxiety in the elevated plus maze, but both knock-out mice showed improved hippocampal-dependent fear context-memory (Fig. 2i,j and Extended Data Fig. 5d-g). While the mice showed no apparent differences in finding the visual platform in the Morris water maze, *Mphosph8*^Δ/Δn^, and to some extent *Morc2a*^Δ/Δn^ mice, performed worse in finding the hidden platform in the Morris water maze test in short-term and long-term probe trials, indicating impaired spatial learning (Extended Data Fig. 5h,i). *Mphosph8* and *Morc2a* knock-out mice did not show any evidence of ataxia or gait-related problems using the catwalk assay (Extended Data Fig. 5j,k). Thus, increased midbrain sizes and accompanying brain architectural changes in *Mphosph8* and *Morc2a* mutant mice are associated with defective motor functions and spatial learning, yet improved fear-context memory.

### Upregulation of clustered protocadherins upon loss of *Mphosph8* or *Morc2a*

Since MPP8 is part of the repressive HUSH complex, we assessed transcriptional changes upon loss of *Mphosph8* and performed RNA-seq from control *Nestin*-Cre+ and *Mphosph8*^Δ/Δn^ littermate whole brains at postnatal day 14. To estimate repetitive element enrichment using the RNA-seq data, we employed the *DESeq2* and *piPipes* pipelines and detected no major changes in repetitive elements, with the exception of a slight upregulation of major satellite repeats (GSAT_MM) and some LTRs and LINE elements (Extended Data Fig. 6a-c). Differentially expressed protein coding genes contained only two downregulated (including *Mphosph8*) and 55 upregulated genes (fold change > 2, adjusted *p* value < 0.05), which were highly enriched in *synapse assembly, synapse organization* and *cell-cell adhesion* gene ontology (GO) categories (Extended Data Fig. 6d). Intriguingly, 21 of the 55 upregulated genes were from the protocadherin (Pcdh) gene cluster on chromosome 18 (Extended Data Fig. 6e). These clustered protocadherins comprise a large family of cell surface molecules expressed in the developing vertebrate nervous system. Pcdh combinations in individual neurons provide a barcode that mediates the precise molecular recognition of synaptic partners and are involved in synapse development^15–17^ and neuronal fidelity. This family of adhesion molecules has also been implicated in brain evolution ^18, 19^ .

Remarkably, the upregulation of protocadherins was detected comparably 2-4-fold in different brain regions upon loss of both *Mphosph8* or *Morc2a* as representatively shown for *Pcdhb14* (Extended Data Fig. 7a,b). Due to the neuron-specific expression of protocadherins, we optimized a protocol for flow cytometry-based sorting of NeuN-immunotagged brain nuclei. Fluorescence-activated cell sorting (FACS) of freshly isolated neuronal nuclei (NeuN+) and non-neuronal nuclei of the brain (NeuN-) revealed an almost exclusive expression of MPP8 and MORC2A in neurons but not in non-neuronal cells of the brain (Fig. 3a,b). Importantly, loss of either *Mphosph8* or *Morc2a* resulted in an upregulation of clustered protocadherins (cPcdhs) in NeuN+ neuronal nuclei (Fig. 3c and Extended Data Fig. 7c). The observed upregulation of cPcdhs could stem from the fact that the expression level of already expressed protocadherins is elevated or that there are increased numbers of protocadherins per cell, i.e. more cells express any particular cPcdh. To explore these hypotheses, we analyzed the expression of a specific protocadherin – *Pcdhb2* – by chromogenic RNA *in situ* hybridization (RNAscope) in *Mphosph8*^Δ/Δn^ and *Morc2a*^Δ/Δn^ brains. We indeed found a higher number and proportion of *Pcdhb2* positive cells compared to wildtype littermates (Extended Data Fig. 7d). To test whether elevated cPcdh expression might expand the neuronal barcodes and increase synaptic connections, we crossed *Morc2a*^Δ/Δn^ animals to mice expressing EGFP under the control of a neuronal promoter (Tg(Thy1-EGFP).MJrs mice), which allowed direct examination of dendritic spines *in vivo* ^20^. We indeed observed increased density of spines in *Morc2a*^Δ/Δn^ brains (Fig. 3d). These data show that in *Mphosph8*^Δ/Δn^ and *Morc2a*^Δ/Δn^ brains individual neurons express more clustered protocadherin isoforms and exhibit a higher number of synaptic connections.

**Figure 3.**
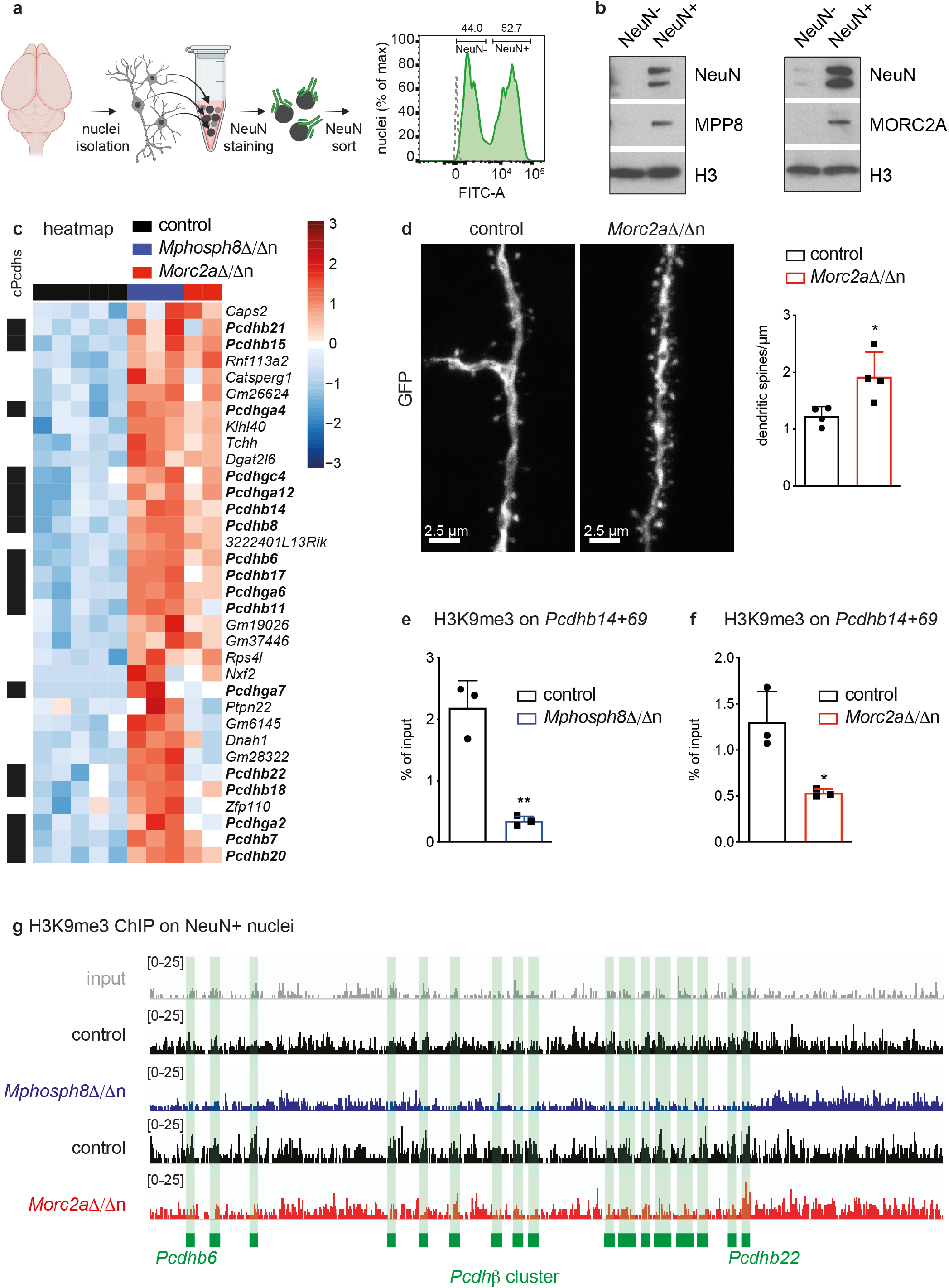
Protocadherin cluster H3K9 trimethylation and synaptic spines in *Mphosph8*- and *Morc2a*-deficient neurons. (**a**) Schematic illustration (created with Biorender.com) and representative plot of flow cytometry-based sorting of NeuN-immunotagged brain nuclei. The gray dotted line in the histogram indicates unstained nuclei as background control. (**b**) Immunoblot analysis of NeuN- and NeuN+ wildtype brain nuclear extracts as depicted in (**a**) using antibodies against MPP8, MORC2A and histone H3 (H3) as loading control. (**c**) Heatmap of upregulated genes (adjusted *p* value < 0.05, fold change > 2 between control and *Mphosph8*^Δ/Δn^) found in an RNA-seq experiment using sorted brain NeuN+ nuclei from control, *Mphosph8*^Δ/Δn^ and *Morc2a*^Δ/Δn^ adult littermates. Note that half of the upregulated genes are from the protocadherin gene cluster (cPcdh, black). Z-score is shown from -3 (blue) to 3 (red). **(d)** (left) Representative maximum intensity projection of z-stacked confocal images of hippocampal CA1 pyramidal cell distal dendrites and (right) quantification of dendritic spines of control versus *Morc2a*^Δ/Δn^ (n = 4) adult littermates (scale bar = 2.5 µm). For each of the 4 mice per genotype the dendritic spine number of 3-6 individual neurons were counted. Data are shown as mean values +/- SD. (**e,f**) Chromatin from control versus *Mphosph8*^Δ/Δn^ littermate (**e**, n = 3) and control versus *Morc2a*^Δ/Δn^ littermate (**f,** n = 3) NeuN+ nuclei was immunoprecipitated with an H3K9me3-specific antibody followed by quantitative reverse transcription polymerase chain reaction (qRT-PCR) with primers specific for *Pcdhb14* (+69 bp from the transcriptional start site). Data are representative for one of two independent experiments and are shown as mean values +/- SD. (**g**) Normalized read density plots of H3K9me3 ChIP-seq in brain NeuN+ nuclei from control, *Mphosph8*^Δ/Δn^ and *Morc2a*^Δ/Δn^ mice. The protocadherin cluster on chromosome 18 is shown, spanning from *Pcdhb6* to *Pcdhb22* (green boxes). For panels d-f each data point represents an individual mouse. *p* values were calculated using the Student’s *t* test. * *p* < 0.05; ** *p* < 0.01.

### *Mphosph8*- and *Morc2a*-deficient neurons show decreased H3K9 trimethylation at the protocadherin cluster

To establish neuronal individuality and generate functional neuronal circuits, in every neuron only a few Pcdh isoforms from the protocadherin gene cluster are transcribed differentially and independently, while the silent protocadherins are marked by DNA^21–23^ methylation and H3K9me3 to maintain the repressed state. To investigate whether the observed upregulation of cPcdhs was associated with decreased trimethylation of H3K9, we first assessed global H3K9me3 levels in *Mphosph8*^Δ/Δn^ brains and did not detect any differences (Extended Data Fig. 7e). When analyzing freshly isolated adult control, *Mphosph8*^Δ/Δn^ and *Morc2a*^Δ/Δn^ NeuN+ neuronal nuclei by chromatin immunoprecipitation (ChIP) experiments for abundance of the histone methylation mark H3K9me3, we however detected markedly decreased H3K9me3 levels specifically at the repetitive-like protocadherin cluster on chromosome 18 (Fig. 3e,f). Chromosome 18 with the clustered protocadherins emerged as the top-scoring locus for H3K9me3 hypomethylation in both *Mphosph8*^Δ/Δn^ and *Morc2a*^Δ/Δn^ neuronal nuclei as visualized in the genome browser views (Fig. 3g) as well as genome-wide Manhattan plots (Extended Data Fig. 7f). Together, this data indicates that *Mphosph8* or *Morc2a* deficiency in neurons leads to decreased histone H3K9 trimethylation specifically on the protocadherin cluster, resulting in upregulation of cPcdhs.

### Human cerebral organoids lacking *MPHOSPH8* or *MORC2* express increased numbers of clustered protocadherins at the single cell level

To translate our mouse data and to model the role of MPHOSPH8 and MORC2 in human brain development, we generated human 3D cerebral organoids ^24, 25^. We established two independent clones of *MPHOSPH8* and *MORC2* knock-out H9 human ES cell lines by CRISPR/Cas9 engineering and used these clones for cerebral organoid generation. Interestingly, we observed an upregulation of MORC2 upon loss of MPHOSPH8 and *vice versa*, suggesting a potential compensation in the mutant ES cell lines (Extended Data Fig. 8a), which to some extent was also detected in *Mphosph8*^Δ/Δn^ and *Morc2a*^Δ/Δn^ mouse brains (Fig. 1b and Extended Data Fig. 2b). We used these ES cell lines to generate cerebral organoids as previously described ^25^. Single cell RNA-sequencing analysis of day 27 cerebral organoids confirmed the enrichment of neuronal progenitors (Extended Data Fig. 8b,c). Fluorescent immunohistochemistry of day 60 cerebral organoids further revealed the presence of both neuronal progenitors and neurons (Extended Data Fig. 8d). *MPHOSPH8* and *MORC2* knock-out human H9 ES cells and cerebral organoids showed a strong upregulation of repetitive elements including LINE-1 and alpha satellites (Extended Data Fig. 8a,e,f), demonstrating that the HUSH complex regulates expression of defined transposons in human stem cells and developing cerebral organoids.

Importantly, while clustered protocadherins are expressed at a very low level in pluripotent human ES cells, an upregulation of clustered protocadherins could be observed in *MPHOSPH8- or MORC2*-deficient cerebral organoids at different time points of differentiation as exemplified by *PCDHB2* (Extended Data Fig. 8g). Single cell RNA-sequencing of day 27 cerebral organoids revealed an increased proportion of cells expressing a defined protocadherin upon loss of *MPHOSPH8* or *MORC2* (Fig 4a,b). The higher number of cells expressing a particular protocadherin was observed throughout all cell clusters including neuronal progenitors (expressing *SOX2*, *PAX6* and *DLX1*) and immature neurons (expressing *DCX*) (Fig. 4a and Extended Data Fig. 8c). Moreover, upon *MPHOSPH8* or *MORC2* knock-out, individual cells expressed a higher number of clustered protocadherins (Fig. 4c-e), especially in the *PCDHB* and *PCDHGA* subclusters (Fig. 4f). Thus, our human cerebral organoid data show that individual *MPHOSPH8- or MORC2*-deficient neurons express increased numbers of clustered protocadherins.

**Figure 4.**
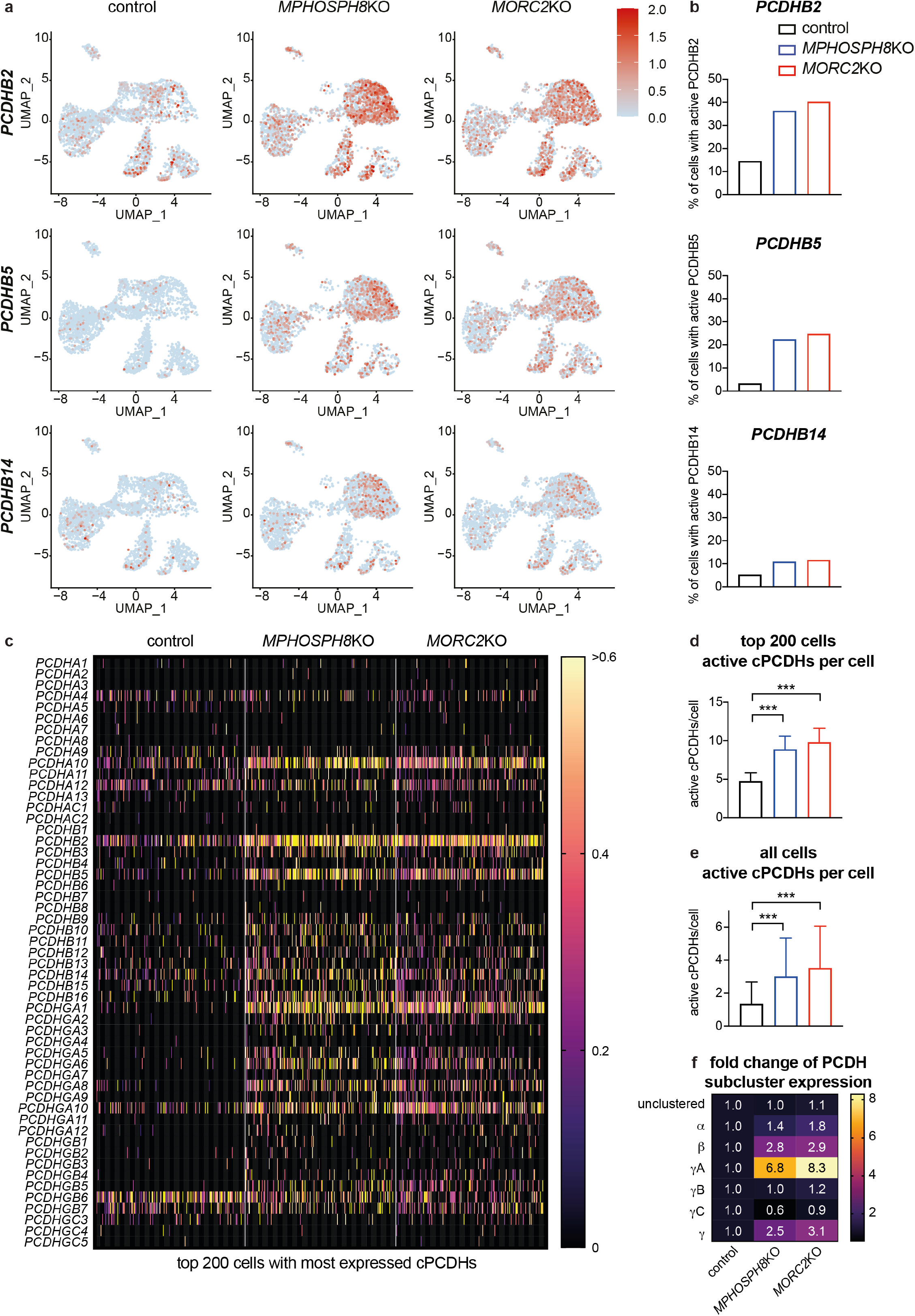
MPHOSPH8 and MORC2 control fidelity of clustered protocadherins in human cerebral organoids. (**a**) Expression of *protocadherin B2* (*PCDHB2*, upper panel), *PCDHB5* (middle panel) and *PCDHB14* (lower panel) in control, *MPHOSPH8*KO and *MORC2*KO single cells from day 27 cerebral organoids projected on a uniform manifold approximation and projection (UMAP) plot. For each group two organoids (each organoid originating from an independent ESC clone) were pooled. Each dot represents one cell and in total approximately 3400 cells per group were analyzed. (**b**) Percent of cells expressing *PCDHB2* (upper panel), *PCDHB5* (middle panel) and *PCDHB14* (lower panel) in control (n = 3488), *MPHOSPH8*KO (n = 3414) and *MORC2*KO (n = 3426) single cells from day 27 cerebral organoids as shown in (**a**). For each group two organoids (each organoid originating from an independent ESC clone) were pooled. (**c**) Heatmap of the normalized expression of the top 200 cells with the most expressed cPCDHs in control, *MPHOSPH8*KO and *MORC2*KO single cells from day 27 cerebral organoids, shown for all clustered protocadherins. For each group two organoids (each organoid originating from an independent ESC clone) were pooled. Log-normalized UMI counts are shown from 0 (black) to > 0.6 (yellow). (**d**) Number of expressed clustered protocadherins per single cell of the top 200 cells with the most expressed cPCDHs in control, *MPHOSPH8*KO and *MORC2*KO single cells from day 27 cerebral organoids as shown in (**c**). Mean values +/- SD are shown. For each group two organoids (each organoid originating from an independent ESC clone) were pooled. (**e**) Number of expressed clustered protocadherins per single cell in control (n = 3488), *MPHOSPH8*KO (n = 3414) and *MORC2*KO (n = 3426) single cells from day 27 cerebral organoids. Mean values +/- SD are shown. For each group two organoids (each organoid originating from an independent ESC clone) were pooled. (**f**) Relative fold increase of expressed unclustered and clustered protocadherins per single cell in *MPHOSPH8*KO and *MORC2*KO single cells from day 27 cerebral organoids in comparison to the control, shown for each subcluster separately. For panels d and e *p* values were calculated using Ordinary one-way ANOVA using multiple comparisons. *** *p* < 0.001.

## Discussion

In this study we identify murine M-phase phosphoprotein 8 (MPP8) and Microrchidia CW-type zinc finger protein 2 (MORC2A) as crucial regulators of brain development and function. Full-body knock-out of either *Mphosph8* or *Morc2a* leads to embryonic lethality and *Nestin-*Cre-mediated nervous system-specific loss of *Mphosph8* or *Morc2a* results in decreased survival and altered brain morphology with expansion of the midbrain leading to increased cerebro-cerebellar distance and more exposed rostral and caudal colliculi. The neuromorphologic and behavioral phenotypes, including motor deficits and differences in learning and memory, are remarkably similar in *Mphosph8* and *Morc2a* knock-out mice. Mechanistically, *Mphosph8* and *Morc2a* are exclusively expressed in neurons, where they repress the protocadherin cluster on mouse chromosome 18 in a H3K9me3-dependent manner, thereby affecting synapse formation.

The tightly controlled stochastic and combinatorial expression of the different protocadherin isoforms in individual neurons might provide the molecular basis for the neuronal diversity, neuronal network complexity and function of the vertebrate brain. In the mouse, the 58 clustered protocadherin genes are tandemly arrayed in α, β, and γ subclusters called *Pcdh*α*, Pcdh*β, and *Pcdh*γ, which encode 14, 22 and 22 cadherin-like proteins, respectively, with each having its own promoter ^26, 27^. The expression of different protocadherin combinations from each of the three gene clusters in individual neurons provides barcodes to distinguish self from non-self and ensures that neurons only interact with other neurons, while they do not contact processes of the same neuron with the same barcode ^28, 29^. By RNAscope RNA *in situ* hybridization and single cell RNA-seq, we observed that individual MPHOSPH8-or MORC2-deficient neurons in both mouse brains and human cerebral organoids express increased numbers of clustered protocadherin isoforms. Importantly, in our mutant mice we also observed increased synapse formation. It remains to be investigated how this is associated with the impaired motor functions, spatial learning deficits and the improved fear-context memory observed in *Mphosph8* or *Morc2a* deficient mice. Since dysregulation of clustered protocadherins is associated with a variety of neurological and neurodevelopmental diseases as well as mental disorders including autism spectrum disorder, bipolar disorder, Alzheimer’s disease, cognitive impairments and schizophrenia ^30, 31^, our data on the key importance of the HUSH complex in protocadherin gene expression might provide new understanding on the epigenetic regulation of such diseases.

The mammalian chromodomain-containing methyl-H3K9 reader protein MPP8 and chromo-like domain-containing MORC2 do not have orthologs either in *Drosophila* or *Caenorhabditis*. This suggests that they could function through different mechanisms than the conserved chromodomain-containing reader protein HP1 and hence represent a second evolutionary route to H3K9me3-mediated heterochromatin regulation in mammalian cells. Interestingly, the upregulated clustered protocadherins are mostly vertebrate-specific and might mediate neurite self-avoidance by specifying single cell identity similar to that of invertebrate Dscam1 proteins. Clustered protocadherins can also be found in some invertebrates such as coleoid cephalopods including octopuses, which have the largest nervous systems among invertebrates and a rich behavioral repertoire including complex problem solving, observational learning and a sophisticated adaptive coloration system ^32^. Remarkably, genomic and transcriptomic sequencing of *Octopus bimaculoides* revealed a high similarity to other invertebrate bilaterians, except for massive expansions in two gene families previously thought to be uniquely enlarged in vertebrates: the protocadherins and the C2H2 superfamily of zinc-finger transcription factors ^18^. In fact, the octopus genome encodes 168 protocadherin genes, nearly three-quarters of which are found in tandem clusters on the genome ^18^. Thus, both octopuses and vertebrates have independently evolved a diverse array of clustered protocadherin genes. The synaptic specificity regulated by members of the protocadherin gene cluster thus allows the enormous increase in number and complexity of neuronal subtypes, synaptic connections and the neural networks of the vertebrate brain compared to most of its invertebrate ancestors. Whether a HUSH-like complex universally controls protocadherin barcoding and regulates brain and cognitive evolution remains to be determined.

Fine-tuning of neural connectivity is important for advanced cerebral functions as well as brain and cognitive evolution. Our data now show the HUSH complex, previously linked to silencing of transposons, has a role in regulating sizes and proportions of brain regions and synapse formation through epigenetic regulation of protocadherin expression.

## Author contributions

J.M.P and A.H. conceptualized, coordinated and designed the experiments. A.H. generated the KO mice and performed experiments including Southern blots, RNA expression, immunoblots, immunohistochemistry, chromatin immunoprecipitations and mouse phenotyping. M.H. counted dendritic spines, assisted in mouse measurements, chromatin immunoprecipitation and quantitative PCR experiments and performed all the cerebral organoid experiments from human ESCs with help from M.B.E. under the supervision of C.E. and J.A.K. A.H. and J.T.-S. optimized and performed the NeuN sort experiments. A.K. coordinated and performed the histology and pathology analyses. M.N. analyzed all the sequencing data. M.O. and D.C. performed PhenoMaster experiments and assisted with organ dissections. U.E. and S.Z. performed teratoma assays and helped with the generation of KO mice. R.K. assisted with mouse genotyping. A.H. and J.M.P. wrote the paper with input from the co-authors.

## Acknowledgments

We thank all members of the Penninger laboratory for helpful discussions and technical support. We are especially thankful to Jelena Zinnanti and Lydia Zopf from the VBCF Preclinical Imaging Facility for their help with magnetic resonance imaging and analysis. Furthermore, we want to thank all members of the IMP-IMBA-GMI Bio-optics Core Facility for assistance in cell/nuclei sorting and imaging as well as the Molecular Biology Service for their help. We also thank the IMBA Stem Cell Core Facility and especially Jana Slovakova for their assistance in H9 cell targeting. Moreover, we thank the Preclinical Phenotyping Facility and especially Sylvia Badurek, the Next Generation Sequencing Facility and the Histopathology Facility at the Vienna Biocenter Core Facilities GmbH (VBCF), member of Vienna Biocenter (VBC), Austria. Furthermore, we thank Hannes Tkadletz from IMP Graphics for his valuable assistance with macroscopic mouse photography and Paul Möseneder for his support in mouse genotyping. J.M.P. is supported by the Austrian Federal Ministry of Education, Science and Research, the Austrian Academy of Sciences and the City of Vienna and grants from the Austrian Science Fund (FWF) Wittgenstein award (Z 271-B19), the T. von Zastrow foundation, and a Canada 150 Research Chairs Program (F18-01336). RIAT-CZ project (ATCZ40) funded via Interreg V-A Austria – Czech Republic is gratefully acknowledged for the financial support of the measurements at the VBCF Preclinical Imaging Facility.

## Methods

### Mice

For *Mphosph8*, the ES cell clone EPD0058_2_C01 (Mphosph8^tm1a(EUCOMM)Wtsi^) was purchased from EUCOMM. For *Morc2a*, the targeting vector PG00072_X_1_H08 (KOMP) was electroporated into IB10/C ES cells generating the Morc2a^tm1a(KOMP)Wtsi^ allele. In both cases exon 4 was flanked by loxP sites and the adjacent LacZ-Neo selection cassette was flanked by Frt sites. Southern blot analysis of ES cells identified correctly targeted clones, which were then used for blastocyst injections to create chimeric mice. These mice were bred on a C57BL/6J genetic background, the LacZ-Neo cassette was removed by crossing mice to an *Actin*-FlpE deleter line (B6.Cg-Tg(ACTFLPe)9205Dym/J) and after various breeding steps homozygous floxed alleles were generated. Then, homozygous floxed *Mphosph8* and *Morc2a* mice were crossed to either *Actin*-Cre (Tmem163^Tg(ACTB-cre)2Mrt^) or *Nestin*-Cre (B6.Cg-Tg(Nes-cre)1Kln/J) mice ^33, 34^. To express EGFP under control of the neuronal *Thy1* promoter and thereby label neurons, mice were crossed to Tg(Thy1-EGFP)MJrs mice ^20^. Mouse genotypes were assessed by PCR (see Extended Data Table 1 for genotyping primer sequences). Of note, only age- and sex-matched littermates from respective crosses were used for experiments. When not explicitly stated, mouse experiments were carried out 10-16 weeks after birth. All mice were bred, maintained, examined and euthanized in accordance with institutional animal care guidelines and ethical animal license protocols approved by the legal authorities. Dorsal and lateral photographs of the head were acquired from selected littermates of each genotype with a Nikon D7 digital camera and a Nikon AF-S Micro Nikkor 105 mm 1:2:8 G VR lens.

### Behavioral experiments

Behavior experiments were carried out under the animal license number GZ:2020-0.392.948 according to Austrian legislation. The open field, Y-maze, elevated plus maze, fear conditioning, grip strength, Morris water maze and CatWalk tests were performed by the Preclinical Phenotyping Facility at the Vienna Biocenter Core Facilities GmbH (VBCF), Vienna Biocenter (VBC), Austria. For testing, mice were transferred to the preclinical phenotyping facility of the VBCF at least 1 week prior to experiments and housed at a 14/10 hours light/dark cycle in IVC racks with access to food and water *ad libitum*. Before each experiment, mice were allowed to habituate to the experimental room for at least 30 minutes prior to any testing.

#### Open field test

Naïve mice were allowed to explore an open field arena (www.tse-systems.com) sized 50 cm (width) x 50 cm (length) x 29.5 cm (height) for 30 minutes with the release from the center and video-tracked using TSE VideoMot 3D version 7.01 software. In the software, a “center” zone was defined as a central square 25 cm x 25 cm in size, the rest being the “border zone”. Light conditions were 200 lux in the center zone. The time spent in each zone, distances travelled, numbers of center visits and rearings were recorded as readout parameters.

#### Y-maze test

The Y-maze was performed as a test for working memory using a custom-built Y-shaped maze with grey, opaque walls and floor with the following dimensions: arm length: 30 cm, arm width: 6 cm, wall height: 14.5 cm. After at least 30 minutes of habituation to the test room (180 lux, visual cues on walls) mice were placed individually into the end of one of the 3 arms (arms A, B, and C), facing the wall at the end of the arm, and allowed to explore the maze for 5 minutes while being video-tracked using the Topscan software (Cleversys Inc, USA). The experimenter watched the videos in the same room, behind a curtain and scored the latency to leave the starting arm (which was alternated between the mice) and the numbers and sequences of arm entries. These sequences were evaluated in terms of the triplet arms. Three arm entries in a row were scored as either correct spontaneous alternations (SA; e.g. BAC, CBA, ABC), erroneous alternate arm returns (AARs; e.g. BAB, CBC, ABA), or erroneous same arm returns (SARs, e.g. BAA, CCB, AAC). After each triplet was scored, the start of the analysis was shifted by one entry and the next triplet sequence was scored. Two such shifts of analyses result in overlapping triplets and the scoring of all possible decision points. Spontaneous alternation performance (SAP) was calculated as [spontaneous alternations (SA)/(total arm entries – 2)].

#### Elevated plus maze test

Mice were placed in the center zone (6.5 x 6.5 cm), facing an open arm of a custom-built elevated plus maze (elevated 54 cm above the floor) with 2 open arms (OA, 30 cm length, 7 cm width) and 2 wall-enclosed arms (closed arms, CA, 30 cm length, 6 cm width, walls 14.5 cm high). Exploration paths were video-tracked for 5 minutes using the Topscan software (Cleversys, Inc., VA, USA) and the amount of time spent and distances travelled in the open arms, closed arms and center zone were recorded. Lux levels were 180 lux in the center zone and open arms and 35 lux in the closed arms.

#### Grip strength test

Grip strength was measured using a grip strength meter (Bioseb, USA). For forelimb measurements, the mouse was gently lowered over the top of a grid so that only its front paws could grip the grid. The animal was gently pulled back and when it released the grid, which is connected to a sensor, the maximal grip strength value of the animal was displayed on the screen and noted. For the forelimb and hindlimb measurements, the mouse was gently lowered over the top of the grid so that both its front and hind paws could grip the grid. The torso was kept parallel to the grid and the mouse was gently pulled back steadily until the grip was released down the complete length of the grid. The maximal grip strength value of the animal was recorded. Both, “forelimb only” and “fore- and hindlimb” tests were performed in an alternating fashion three times/mouse with 15 minutes inter-trial intervals and the values averaged among the three trials.

#### Morris water maze

Mice were trained to swim in a pool (diameter: 1 m) with colored water (white paint used: OBI PU Buntlack schneeweiß glänzend 17.05.23) treated with aquarium cleaner (TetraAqua aquarium conditioner, according to manufacturer’s instructions) at a temperature of 20-22°C. The platform (diameter: 10 cm) was submerged about 8 mm under the water surface, using the visual cues placed in the room for orientation. Mice were placed in the water facing the wall of the pool and given 2 sessions a day with 4 trials per session using alternating entry points in different quadrants for each trial and video-tracked using the software Topscan 3.0 (Cleversys Inc., VA, USA). On day 1, the visual capacity of the mice was assessed by making the platform visible with a black flag and letting the mice explore the pool for 1 minute or until they found the platform. If they did not find the platform within 1 minute of the first session, they were guided towards the platform by pointing a pair of forceps in front of their nose. After a break on days 2 and 3, mice were given hidden platform training from days 4-8 without any marking of the platform. The time needed to find the platform was recorded. After the last trial on day 8, mice were tested for short term memory by removing the platform and letting the mice explore the pool for 1 minute. This probe trial was repeated in the morning of day 11 to test for long term memory. The time spent searching in the target quadrant and target zone (exact location of the platform) was evaluated.

#### Fear conditioning

In the morning of day 1 (09:00am to 01:00pm) mice were trained to associate the conditioned sound stimulus (CS = 85 dB, 10 kHz) to an unconditioned foot shock stimulus (US = 0.5 mA) using the following protocol: 120 sec pre-phase (no sound, no shock), 30 sec CS with the last 2 sec coupled to the US, 90 sec inter-tone-interval (no sound, no shock), 30 sec CS with the last 2 sec coupled to the US, 70 sec post phase (no sound, no shock) in a Coulbourn Habitest operant cage (Coulbourn Instruments, MA, USA, www.coulbourn.com) using house light in the visible range. Mice were videorecorded using the software FreezeFrame from Actimetrics, IL, USA (www.actimetrics.com). For context testing, mice were placed back in the same conditioning box 24 hours later and observed for 4 minutes without any sound or shock presentation. For the cue test, mice were placed in the same fear conditioning box, but with changed appearance (different texture of walls and floor, infrared house light, no ventilator, lemon aroma spotted in box) for approx. 28 hours (in the afternoon session 01:00pm to 17:00pm) after the context test and the following protocol was applied: 120 sec pre phase (no sound, no shock), 60 sec CS (only sound, no shock), 60 sec inter-tone-interval, 60 sec CS (only sound, no shock), 60 sec post-phase (no sound, no shock). Freezing was defined as a minimum of 2 sec without movement except breathing and was analyzed using the FreezeView software from Actimetrics, IL, USA (www.actimetrics.com).

#### CatWalk test

Gait was analyzed using the CatWalk XT system (Noldus Information Technology) essentially as described in ref. ^35^ with the following exceptions: Mice were not trained to reach a goal box, but to directly reach the homecage. Training was performed on the first day and the test was done on two consecutive days directly after the training day. An average of three trials performed on each test day was calculated for each mouse, then an average across the two test days was calculated for each animal.

#### Motor activity in PhenoMaster cages

Activity measurements were performed at room temperature (21°C-23°C) on a 12/12 hours light/dark cycle in a PhenoMaster System (TSE systems, Bad Homburg, Germany) using an open circuit calorimetry system and animals were checked daily by veterinary staff. Mice were housed individually, trained on drinking nozzles for 72 hours and allowed to adapt to the PhenoMaster cage for 2 days. Food and water were provided *ad libitum* in the appropriate devices and measured by the built-in automated instruments. Activity parameters were measured for 5 consecutive days.

#### Accelerating Rotarod

Up to 4 mice were placed simultaneously on the rod compartments of a Rotarod device (Ugo Basile) to evaluate the motor ability. The assays were conducted in the afternoon, between 02:00pm to 5:00pm. One day before the test sessions, the mice were trained to stay on the Rotarod apparatus with a rotation of 5 rpm for 5 minutes as a habituation trial. From day 1 to day 4, the mice were assayed with accelerating mode from 5 to 40 rpm for 5 minute time periods each. The latencies until the mice fell were recorded and passive rotations (clinging onto the rod without running) were treated like a fall.

### Teratoma formation

For teratoma formation, 1 million ES cells (Haplobank) were mixed with Matrigel (Corning, 356231) and injected subcutaneously into Crl:NU(NCr)-Foxn1 nude mice. Teratoma growth was monitored for 4-6 weeks.

### Histology, immunostaining and *in situ* hybridization

Mouse brains were harvested, macroscopically inspected, and fixed overnight by immersion in 4% paraformaldehyde, processed with an automated tissue processor (Logos, Milestone Medical or Donatello, DiaPath/Sanova), embedded in paraffin, and sectioned at a thickness of 2 µm with a standard rotary microtome (Microm HM 355, Thermo Scientific). Sections were stained with Hematoxylin and Eosin (H&E) on an automated staining platform (HMS 740, Microm or Gemini AS, Thermo Scientific) or with Luxol Fast Blue - Cresyl Violet (LFBCV) using a standard manual protocol. Immunohistochemistry (IHC) for the Neuronal Nuclear Antigen (NeuN) was performed using an automated immunostainer (Bond III, Leica or Infinity i6000, Biogenex). Slides were rehydrated, subjected to heat induced antigen retrieval in a citrate buffer and incubated with a mouse anti-NeuN antibody (clone A60, Millipore, MAB377, 1:100). The used secondary antibodies were mouse linker antibody (Abcam, ab133469) and a goat anti-rabbit antibody (Dako, E0432, 1:500). Chromogenic detection was performed with the Bond Intense R Detection System (Leica, DS9263) or the DCS Supervision 2 Polymer System (Innovative Diagnostik-Systeme, PD000POL-K) and the DAB substrate kit (Abcam, ab64238). *In situ* hybridization for Protocadherin beta 2 (*Pcdhb2*) was performed with a manual protocol using the RNAscope® Probe- Mm-Pcdhb2 (ACD, 46781). Chromogenic detection was performed using the RNAscope® 2.5 HD Reagent Kit - RED (ACD).

Slides were reviewed by a board-certified pathologist with an Axioskop 2 MOT microscope (Zeiss). Whole slide images were prepared using the Pannoramic FLASH 250 III whole slide scanner (3D Histech) with the 40x/0.95 plan apochromat objective and Adimec Quartz Q12A180 camera. Representative images were acquired from slides with a SPOT Insight 2 camera (Spot Imaging, Diagnostic Instruments) and from whole slide images with the Case Viewer Software (3D Histech). Quantification of NeuN positive cells was performed with QuPath, an open source whole slide image analysis platform ^36^. For quantification of positive cells in the cortex, diencephalon, tectum (rostral and caudal colliculi), and tegmentum, these regions were manually annotated in each image within each workspace. For quantification of positive cells in the cerebellum, the granular layer and other regions (molecular layer and white matter) were separately delineated with the ANN_MLP Pixel classifier. Positive cells were detected by setting the intensity thresholds for the nuclear DAB OD mean score compartment. The threshold was applied to all images within the workspace with a command history script. Detection was performed with separate thresholds and scripts for 1) the granular layer of the cerebellum, 2) the molecular and white matter layers of the cerebellum, and 3) all the other indicated regions of the brain. Examples of the regions and corresponding cellular detections are presented in Extended Data Fig. 3c,d. Quantification of *Pcdhb2* positive spots was performed by manual enumeration of positive spots and nuclei in the cerebellar granular and molecular layers using FIJI ^37^ with exported TIFF files from whole slide images.

### qRT-PCR and qRT-PCR analysis

Tissues and cells were isolated and homogenized in TRIzol reagent (Invitrogen). Total RNA was isolated according to the manufacturer’s instructions. RNA was reverse transcribed with the iScript cDNA synthesis kit (Bio-Rad). Real-time PCR analysis was performed with GoTaq qPCR master mix (Promega) on a CFX384 system (Bio-Rad). Data were normalized to values for the housekeeping gene *Gapdh* for mouse samples and *ACTB* (encoding ß-Actin) for human samples. All primers used in this study are listed in Extended Data Table 1.

### mRNA sequencing and data analysis

cDNA libraries for each of the samples were generated from total RNA using the NEB PolyA enrichment kit following the manufacturer’s instructions. SR50 Sequencing was performed on HiSeq 2500 at the VBCF NGS Unit (www.viennabiocenter.org/facilities) using v4 SBS-reagents. RNA-seq reads were trimmed using trim-galore v0.5.0 and reads mapping to abundant sequences included in the iGenomes UCSC GRCm38 reference (mouse rDNA, mouse mitochondrial chromosome, phiX174 genome, adapter) were removed using bowtie2 v2.3.4.1 alignment. Remaining reads were aligned to the mouse genome (Ensembl GRCm38 release 94) using star v2.6.0c and reads in genes were counted with featureCounts (subread v1.6.2) and TEcounts v2.0.3. Differential gene expression analysis on raw counts of genes or genes plus repeat elements was performed using DESeq2 v1.18.1, and gene set over-representation analysis was performed with clusterprofiler 3.6.0 in R v3.4.1. The data discussed in this publication have been deposited in NCBI’s Gene Expression Omnibus ^38^ and are accessible through the GEO Series accession number GSE185330.

### Protein isolation and immunoblotting

For protein extraction, cells or tissues were manually homogenized in Hunt buffer (20 mM Tris-HCl pH 8.0, 100 mM sodium chloride, 1 mM EDTA, 0.5% NP-40) supplemented with Halt protease/phosphatase inhibitor cocktail (Thermo Scientific). After full-speed centrifugation, the supernatant containing the soluble protein fraction was further used. Equal amounts of 20 to 30 μg of protein were separated by SDS-PAGE and transferred onto PVDF membranes (Immobilion-P, Merck Millipore) according to standard protocols. Blots were blocked for 1 hour with 5% milk in TBST (1x TBS and 0.1% Tween-20) and were then incubated overnight at 4°C with primary antibodies diluted in 5% milk in TBST. Blots were washed 3 times in TBST for 5 minutes and further incubated with HRP-conjugated secondary anti-mouse-IgG-H&L chain (Promega) or anti-rabbit-IgG-F(ab’)2 (GE Healthcare) antibody for 1 hour at room temperature, washed 3 times in TBST for 5 minutes and visualized using enhanced chemiluminescence (ECL, GE Healthcare). See Extended Data Table 1 for a list of antibodies used in this study. β-Actin was used to control for protein loading.

### Sorting of neuronal nuclei

Freshly isolated mouse brains were homogenized in nuclear isolation buffer (10 mM Tris-HCl, 50 mM sodium-disulfite, 1% Triton X-100, 10 mM MgCl_2_, 8.6% sucrose, pH 6.5) and centrifuged 5 minutes at 455 g at 4°C. The resulting pellet was washed 4 times with 900 µl nuclear isolation buffer with centrifugation at 455 g at 4°C. The pelleted nuclei were then resuspended in 1.4 ml of PBS, filtered through a cell strainer and stained with a 1:1000 dilution of the anti-NeuN antibody (clone A60, Alexa Fluor 488 conjugated, Millipore MAB377X) for 45 minutes by rotation in the dark at 4°C after adding BSA to a final concentration of 0.1%. After washing in PBS, the stained nuclei were filtered through FACS tubes, sorted on a FACS Aria III (BD) sorter in PBS, pelleted 5 minutes at 590 g at 4°C and the pellet was immediately used for RNA isolation or frozen at -80°C.

### Chromatin immunoprecipitation (ChIP)

Isolated brains were finely minced and homogenized in PBS, washed with PBS and cross-linked by adding formaldehyde (to a final concentration of 1%) at room temperature for 10 minutes. The cross-linking process was stopped by addition of glycine to a final concentration of 125 mM. The chromatin isolation procedure was followed as previously described ^39^. For ChIP assays, equal amounts of Bioruptor-sonicated chromatin were diluted 10-fold and precipitated overnight with the following antibodies: H3K9me3 (Abcam) and rabbit IgG (Invitrogen) as a control. Chromatin antibody complexes were isolated using protein A-beads for rabbit primary antibodies or G-beads for mouse primary antibodies (Dynabeads, Invitrogen). The PCRs with 1:20 dilutions of genomic DNA (input) were carried out together with the precipitated DNA. The extracted DNA was used for quantitative PCR analysis using the primers listed in Extended Data Table 1 and also for ChIP-seq. For neuron-specific ChIP, freshly isolated brains were homogenized, cross-linked, treated with glycine, washed and used for FACS sorting of neuronal brain nuclei (NeuN+) as described above. After the FACS-sort, the pelleted nuclei were immediately resuspended in ChIP lysis buffer (50 mM Tris-HCl, 10 mM EDTA, 1% SDS, pH 8.1) and continued as described above.

### Deletion of MPHOSPH8 and MORC2 in human embryonic stem cells

The human embryonic stem cell (hESCs) line WA09 (H9) was obtained from WiCell, verified to display a normal karyotype. H9 cells were cultured feeder-free on hESCs-qualified Matrigel (Corning, 354230)-coated plates in E8 Medium (IMBA Stem Cell Core Facility) in DMEM/F-12 (Gibco, 11320033) with an Antibiotic-Antimycotic cocktail (Thermo Fisher, 15240062). All stem cells were maintained in a 5% CO_2_ incubator at 37°C. Cells were split using 0.5 mM UltraPure EDTA (Invitrogen, 15575-038) in PBS to wash once, then incubated for 3 minutes, EDTA aspirated, rinsed in E8 Medium and plated on Matrigel-coated plates. For the generation of knock-out cells, the pSpCas9(BB)-2A-Puro (PX459: Addgene, 62988) plasmid was used carrying a gRNA targeting *MPHOSPH8* (clone C7 and C11: gCATAGACGATCACAAAACCA) or *MORC2* (clone A6 and C4: gACCAAACAAGAATTCGTGAG). H9 cells were nucleofected using the Amaxa nucleofector (Lonza) with the P3 primary cell 4D nucleofector kit L (Lonza LONV4XP-3024). Puromycin (Invivogen 10 mg/ml, ant-pr-1) selection with 0.2 µg/ml in E8 medium started two days after transfection. Puromycin was removed after two days and the cells were further kept in E8 medium. After selecting bulk clones by TIDE analysis (http://shinyapps.datacurators.nl/tide), single cell colonies were picked and verified to carry frame-shift mutations. Loss of MPHOSPH8 and MORC2 protein in hESCs was confirmed by immunoblots.

### Human cerebral organoids

Cerebral organoids were generated and cultured as previously described ^25^ with slight modifications. Briefly, H9 hESC cells were grown to 60-80% confluency, washed with PBS and single cell suspensions were obtained using Accutase (Sigma, A6964). Pelleted cells were resuspended in E8 media supplemented with RevitaCell cell supplement (Invitrogen, A2644501) and counted. 9000 cells were seeded per well to form embryoid bodies in a 96-well ultra-low-attachment U-bottom plate (Sbio, MS-9096UZ) in 150 μl E8 medium supplemented with RevitaCell. On day 3, media was changed to E8, and from day 6 to day 13 organoids were grown in neural induction media (∼1 L neural induction media containing 1 L DMEM/F-12, 10 ml N2 supplement (IMBA Molecular Biology Service), 10 ml GlutaMAX (Invitrogen, 35050-038), 10 ml MEM-NEAA (Sigma, M7145), 1 ml Heparin (Sigma, H3149) of a 1 mg/ml dilution), slightly different as previously described ^25, 40^. On day 10, embryoid bodies were embedded into Matrigel (Corning, 354234) droplets, transferred to a 10 cm dish and cultured from day 13 until day 25 in differentiation media without vitamin A (∼1 L differentiation media containing 500 ml DMEM/F-12 and 500 ml neurobasal media (Invitrogen, 21103049), 5 ml N2 supplement, 10 ml GlutaMAX, 5 ml MEM-NEAA, 20 ml B27 supplement minus vitamin A (Invitrogen, 12587010), 350 µl of a 1:100 dilution of 2-Mercaptoethanol (Merck, 8057400250), 250 µl insulin (Sigma, I9278-5ML)), afterwards in differentiation medium with vitamin A (IMBA Molecular Biology Service), 0.4 mM vitamin C (Sigma, 255564-100G) and 0.1% m/v NaHCO_3_ (Merck, 1.06329.1000). From day 20 onwards, cerebral organoids were constantly shaken on a horizontal shaker.

### Single cell RNA-sequencing of cerebral organoids

Single cell suspensions from two cerebral organoids per condition were prepared at day 27 using 0.9x:1x Accutase/Trypsin (Gibco, 11538876) and incubated for 15 minutes on 37°C while shaking and dissociating manually by pipetting every 5 minutes. Digestion was stopped by adding gradually two volumes of cold DMEM/F-12, washed with DPBS -/- (Thermo Fisher, 14190094) plus 0.04% BSA (VWR, 9048-46-8). Cells were then filtered through a cell strainer, counted and loaded onto a 10x Genomics controller in DPBS -/- plus 0.04% BSA according to the manufacturer’s protocol with a target cell number of 4000 cells. Sequencing was performed on an Illumina NextSeq2000 lane and analyzed using 10x Cell Ranger 4.0.0 workflow, reference refdata-gex-GRCh38-2020-A and Seurat 3.2.2.

### Immunohistochemistry of cerebral organoids

Organoids were washed 3 times in PBS and fixed overnight at 4°C in 4% paraformaldehyde. The next day, the organoids were washed again 3 times in PBS and dehydrated overnight at 4°C in 30% sucrose. The organoids were then embedded with OCT in a multiwell format and stored at -80°C until cryo-sectioning. The block was cut in 20 mm sections and the sections were stored at -80°C. For immunostaining, the slides were dried at room temperature, washed 3 times in PBS and blocked in 4% BSA, 0.5% Triton X-100 in PBS. Afterwards the sections were incubated with the primary antibody overnight in a humidified chamber. The next day, the sections were washed 3 times in washing solution (PBS containing 0.05% Triton X-100) and incubated for 2 hours at room temperature in the secondary antibody and DAPI (Invitrogen, D3571, dilution 1:1000). The slides were afterwards washed 3 times in washing solution, mounted and sealed. See Extended Data Table 1 for a list of primary and secondary antibodies used for immunostaining of the cerebral organoids.

### Statistics

All values are given as means ± standard deviations (SD) unless stated otherwise (SEM in behavior experiments in Fig. 2 and Extended Data Fig. 5). Comparisons between two groups were analyzed using unpaired Student’s *t* tests and corrected for multiple comparisons using the Holm-Sidak method. Comparisons between more than two groups were analyzed using Ordinary one-way ANOVA (Fig. 4d,e and Extended Data Fig. 8e-g). Survival curves (Fig. 1d, Extended Data Fig. 2d and Extended Data Fig. 4i) were compared by the Log-rank (Mantel-Cox) test. *p* values were calculated with GraphPad Prism software: *, *p* < 0.05; **, *p* < 0.01; ***, *p* < 0.001; ns, not significant.

## Extended Data Figure legends

**Extended Data Figure 1. Generation of conditional *Mphosph8* or *Morc2a* knock-out ES cells and mice.**

(**a,b**) Schematic illustration of the conditional *Mphosph8* (**a**) and *Morc2a* (**b**) alleles. Exons are depicted as black numbered boxes and Frt and loxP sites are shown as green and red triangles, respectively. The LacZ-Neo cassette can be removed by crossing mice to a *FlpE* deleter line. LoxP flanked exon 4 can be removed upon recombination in mice expressing Cre recombinase. En2 SA = mouse En2 splicing acceptor acceptor, T2A = T2A element, pA = polyadenylation signal, IRES = internal ribosome entry side, hbactP = human beta actin promoter. (**c,d**) Southern blot analysis of control versus heterozygous *Mphosph8*^fl/+^ (**c**) and control versus heterozygous *Morc2a*^fl/+^ (**d**) ES cells indicating correct targeting. (**e,f**) Teratoma formation 4 weeks after injection of control versus *Mphosph8* knock-out ES cells (**e**) and control versus *Morc2a* knock-out ES cells (**f**). (**g,h**) Genotype statistics of born mice with *Actin-*Cre-mediated full-body knock-out of *Mphosph8* (**g**, n = 39) and *Morc2a* (**h**, n = 59).

**(i)** Genotype statistics of embryonic day E11.5 mice with *Actin-*Cre-mediated full-body knock-out of *Mphosph8* (n = 19).

**(j)** Representative photographs of control (left) and *Actin-*Cre-mediated full-body knock-out of *Morc2a* (right) embryos at embryonic day E11.5 (upper panel) and E13.5 (lower panel). (**k,l**) Immunoblot analysis of adult wildtype tissue extracts with antibodies against MPP8 (**k**), MORC2A (**l**) and β-Actin as loading control.

**Extended Data** **Figure 2. Altered brain architecture in *Mphosph8* knock-out mice.**

**(a)** Relative mRNA expression of *Mphosph8* in littermate control and homozygous *Mphosph8*^Δ/Δn^ brains. Mean values +/- SD were normalized to the housekeeping gene *Gapdh* (n = 3). Data are shown for one of two independent experiments.

**(b)** Immunoblot analysis of littermate control versus *Mphosph8*^Δ/Δn^ whole brain extracts using antibodies against MPP8, MORC2A and β-Actin as loading control. The asterisk indicates an unspecific band.

**(c)** Body weights of *Mphosph8*^fl/fl^ (n = 18), *Nestin-*Cre+ (n = 8), heterozygous *Mphosph8*^Δ/+n^ (n = 7) and *Mphosph8*^Δ/Δn^ (n = 9) male littermates +/- SD during the first 3 months of age. Data from several litters were pooled.

**(d)** Kaplan-Meier survival curve over the first 300 days of control versus *Mphosph8*^Δ/Δn^ littermates. Data from several litters were pooled (n > 27) and survival curves were compared by the Log-rank (Mantel-Cox) test.

**(e)** Representative macroscopic photographs of control (left) and *Mphosph8*^Δ/Δn^ (right) mouse heads (scale bar = 1 cm).

**(f)** Body weights, brain weights and brain/body ratios of control compared to *Mphosph8*^Δ/Δn^ adult (at 3-4 months) littermates. Data from two independent experiments were pooled and shown as mean values +/- SD (n = 8).

**(g)** Representative macroscopic images of a control (left) and an *Mphosph8*^Δ/Δn^ (right) littermate adult (at 3-4 months) brain.

**(h)** Representative *in vivo* MRI brain scans of control (left) and *Mphosph8*^Δ/Δn^ (right) adult (at 3-4 months) littermates (scale bar = 10 mm). In total 3 mice per genotype were analyzed.

**(i)** NeuN immunohistochemistry (IHC) in adult (at 3-4 months) control (left) and *Mphosph8*^Δ/Δn^ (right) brains (scale bar = 2 mm). In total at least 3 mice per genotype were analyzed by immunohistochemistry. OB = olfactory bulb, CTX = cerebral cortex, HC = hippocampus, RC = rostral colliculus, CC = caudal colliculus, TM = tegmentum, CB = cerebellum. Arrows indicate widening of collicular regions of the midbrain in *Mphosph8*^Δ/Δn^ animals. For panels a and f each data point represents an individual mouse. *p* values were calculated using the Student’s *t* test. ** *p* < 0.01; *** *p* < 0.001; ns, not significant.

**Extended Data Figure 3. Increased density of NeuN+ neurons in defined brain regions of *Mphosph8*- or *Morc2a*-deficient mice. (a,b) Quantification of NeuN positive neurons in six brain regions of *Mphosph8*^Δ/Δn^**

**(a)** and *Morc2a*^Δ/Δn^ mice (**b**). For rostral colliculus, caudal colliculus, cerebral cortex and thalamus data from two independent experiments were pooled and shown as mean values +/- SD (n > 4). For the tegmentum and the cerebellum (granular layer) data from one experiment are shown as mean values +/- SD (n = 3). (**c,d**) Representative immunodetection of NeuN positive neurons in *Mphosph8*^Δ/Δn^ (**c**) and *Morc2a*^Δ/Δn^ (**d**) mice using QuPath v0.2.3 (scale bar = 1 mm). Five brain regions are marked, including the rostral colliculus (RC), caudal colliculus (CC), cerebral cortex (CTX), thalamus (TH) and tegmentum (TM) (left) and the granular layer of the cerebellum (CB) (right). (**e,g**) Histomorphologic features of the hippocampal formation in *Mphosph8*^Δ/Δn^ (**e**) and *Morc2a*^Δ/Δn^ brains (**g**) stained with Luxol Fast Blue -Cresyl Violet (LFBCV).

(**f,h**) Histomorphologic features of the cerebral cortical layers in *Mphosph8*^Δ/Δn^ (**f**) and *Morc2a*^Δ/Δn^ brains (**h**) stained with Luxol Fast Blue - Cresyl Violet (LFBCV). For panels a and b each data point represents an individual mouse. *p* values were calculated using the Student’s *t* test. * *p* < 0.05; ** *p* < 0.01; *** *p* < 0.001; ns, not significant.

**Extended Data Figure 4. *Mphosph8*^Δ/Δn^ *Morc2a*^Δ/Δn^ double knock-out mice.**

**(a)** Relative mRNA expression of *Mphosph8* (left) and *Morc2a* (right) in littermate control and homozygous *Mphosph8*^Δ/Δn^ *Morc2a*^Δ/Δn^ double mutant brains. Mean values +/- SD were normalized to the housekeeping gene *Gapdh* (n > 3). Data are shown for one of two independent experiments.

**(b)** Immunoblot analysis of littermate control versus *Mphosph8*^Δ/Δn^ *Morc2a*^Δ/Δn^ brain extracts with antibodies against MPP8, MORC2A and β-Actin as loading control. The asterisk indicates an unspecific band.

**(c)** Body weights of control *Mphosph8*^fl/fl^ *Morc2a*^fl/fl^ (n = 18), *Nestin-*Cre+ (n = 8) and *Mphosph8*^Δ/Δn^ *Morc2a*^Δ/Δn^ (n = 14) male littermates +/- SD during the first 3 months of age. Data from several litters were pooled.

**(d)** Representative macroscopic photographs of control (left) and *Mphosph8*^Δ/Δn^ *Morc2a*^Δ/Δn^ (right) mouse heads (scale bar = 1 cm).

**(e)** Body weights, brain weights and brain/body ratios of control compared to *Mphosph8*^Δ/Δn^ *Morc2a*^Δ/Δn^ adult (at 3-4 months) littermates. Data from one experiment are shown as mean values +/- SD (n = 4).

**(f)** Representative macroscopic images of a control (left) and a *Mphosph8*^Δ/Δn^ *Morc2a*^Δ/Δn^ (right) littermate adult (at 3-4 months) brain.

**(g)** Representative *in vivo* MRI brain scan of control (left) and *Mphosph8*^Δ/Δn^ *Morc2a*^Δ/Δn^ (right) adult littermates (scale bar = 10 mm). In total 3 mice per genotype were analyzed.

**(h)** NeuN immunohistochemistry (IHC) in adult (at 3-4 months) control (left) and *Mphosph8*^Δ/Δn^ *Morc2a*^Δ/Δn^ (right) brains (scale bar = 2 mm). At least 3 mice per genotype were analyzed by immunohistochemistry. OB = olfactory bulb, CTX = cerebral cortex, HC = hippocampus, RC = rostral colliculus, CC = caudal colliculus, TM = tegmentum, CB = cerebellum. Arrows indicate widening of collicular regions of the midbrain in *Mphosph8*^Δ/Δn^ *Morc2a*^Δ/Δn^ brains.

**(i)** Kaplan-Meier survival curve over the first 300 days of control versus *Mphosph8*^Δ/Δn^ *Morc2a*^Δ/Δn^ littermates. Data from several litters were pooled (n > 18) and survival curves were compared by the Log-rank (Mantel-Cox) test. For panels a and e each data point represents an individual mouse. *p* values were calculated using the Student’s *t* test. ** *p* < 0.01; *** *p* < 0.001; ns, not significant.

**Extended Data Figure 5. Behavioral assessment of *Mphosph8* or *Morc2a* mutant mice.**

**(a)** Daily activity in PhenoMaster cages measured by beam breaks of *Nestin-*Cre*+* mice compared to *Nestin-*Cre- littermates over 24 hours. Data from two independent experiments were pooled and shown as mean values +/- SEM (n > 5).

**(b)** Global distances travelled and numbers of rearings of *Nestin-*Cre*+* mice compared to *Nestin-*Cre- littermates in the open field test. Data from two independent experiments were pooled and shown as mean values +/- SEM (n > 5).

**(c)** Latency to fall from the accelerating Rotarod of *Nestin-*Cre*+* mice compared to *Nestin-*Cre- littermates. Data from two independent experiments were pooled and shown as mean values +/- SEM (n > 5). (**d,e**) Spontaneous alteration performance (SAP) in the Y maze of *Mphosph8*^Δ/Δn^ mice (**d**) and *Morc2a*^Δ/Δn^ mice (**e**) compared to their respective control littermates. Data are shown as mean values +/- SEM (n > 7, two independent experiments are pooled). (**f,g**) Percent time spent in the three compartments of the elevated plus maze by *Mphosph8*^Δ/Δn^ mice (**f**) and *Morc2a*^Δ/Δn^ mice (**g**) compared to control littermates. Data are representative of one of two independent experiments and shown as mean values +/- SEM (n > 4). (**h,i**) Latency to find the visual platform in the Morris water maze (left panels) and latency to find the hidden platform in the Morris water maze in the short-term (ST-PT) and long-term (LT-PT) probe trial (right) of *Mphosph8*^Δ/Δn^ mice (**h**) and *Morc2a*^Δ/Δn^ mice (**I**) compared to control littermates. Data are representative of one of two independent experiments and shown as mean values +/-SEM (n > 4).

(**j,k**) Average speed in the Noldus CatWalk system of *Mphosph8*^Δ/Δn^ mice (**j**, n = 4) and *Morc2a*^Δ/Δn^ mice (**k**, n = 6) compared to control littermates. Data are shown as mean values +/- SEM. For panel a (right part) and panels d-k each data point represents an individual mouse. *p* values were calculated using the Student’s *t* test. * *p* < 0.05; ** *p* < 0.01; ns, not significant.

**Extended Data Figure 6. Upregulation of the protocadherin cluster in the brains of *Mphosph8* knock-out mice.**

**(a)** Volcano plot of differential expression analysis (*DESeq2*) to estimate repetitive element enrichment in an RNA-seq experiment of 3 littermate control and 3 *Mphosph8*^Δ/Δn^ brains on postnatal day 14. Elements with a fold change > 2 (log2 fold change > 1) are shown in red, others in gray. GSAT_MM = major satellite repeats, MMERGLN-int:ERV1:LTR = LTR retrotransposon belonging to the ERV1 family.

**(b)** *piPipes* analysis to estimate repetitive element enrichment in an RNA-seq experiment of 3 littermate control and 3 *Mphosph8*^Δ/Δn^ brains on postnatal day 14. NM = mRNA, NR = ncRNA. GSAT_MM = major satellite repeats.

**(c)** Relative mRNA expression of major satellites repeats (GSAT) in littermate control and *Mphosph8*^Δ/Δn^ brains. Mean values +/- SD were normalized to the housekeeping gene *Gapdh* (n = 3).

**(d)** Biological gene ontology (GO) processes overrepresented among the gene set of 55 upregulated genes in an RNA-seq experiment of 3 littermate control and 3 *Mphosph8*^Δ/Δn^ brains on postnatal day 14 (adjusted *p* value < 0.05). The negative log10 of the q-value to control the positive false discovery rate for multiple testing is shown.

**(e)** Heatmap of differentially regulated genes (adjusted *p* value < 0.05, fold change >2) found in the RNA-seq experiment illustrated in **(d)** showing approximately half of the upregulated genes are from the protocadherin gene cluster (cPcdh, marked in black). Z-score is shown from -3 (blue) to 3 (red). For panel c each data point represents an individual mouse. *p* values were calculated using the Student’s *t* test. * *p* < 0.05.

**Extended Data Figure 7. Loss of murine *Mphosph8* or *Morc2a* leads to upregulation of clustered protocadherins.**

(**a,b**) Relative mRNA expression of *Pcdhb14* in littermate control versus *Mphosph8*^Δ/Δn^ (**a**) and control versus *Morc2a*^Δ/Δn^ (**b**) brain areas including the olfactory bulb, cortex, midbrain, pons/medulla and cerebellum. Mean values +/- SD were normalized to the housekeeping gene *Gapdh* (n = 3) and the control values were set to 1.

(**c**) Relative mRNA expression of *Mphosph8*, *Morc2a* and *Pcdhb14* in littermate control, *Mphosph8*^Δ/Δn^ and *Morc2a*^Δ/Δn^ sorted brain NeuN+ nuclei. Mean values +/- SD were normalized to the housekeeping gene *Gapdh* (n = 2).

(**d**) *In situ* hybridization (RNAscope) (left) for *Pcdhb2* mRNA and quantification (right panels) in the granular cell layer of the cerebellum in control versus *Mphosph8*^Δ/Δn^ (upper panel, n = 4) and control versus *Morc2a*^Δ/Δn^ (lower panel, n = 3) adult (at 3-4 months) littermates (scale bar = 50 µm). Data are shown as mean values +/- SD.

(**e**) Immunoblot analysis of littermate control, heterozygous *Mphosph8*^Δ/+n^ versus *Mphosph8*^Δ/Δn^ brain extracts with antibodies against MPP8 and H3K9me3 as well as β-Actin and total H3 as loading controls.

(**f**) Manhattan plot visualization of H3K9me3 ChIP-seq differential analysis using diffreps in 1kb sliding windows showing localized enrichments for decreased abundance of H3K9me3 methylation in NeuN+ nuclei of *Mphosph8*^Δ/Δn^ versus control mice (left, n = 3) and *Morc2a*^Δ/Δn^ versus control mice (right, n = 2). All the top-scoring dots on chromosome 18 correspond to the clustered Pcdh locus. For panels a-d each data point represents an individual mouse. *p* values were calculated using the Student’s *t* test. * *p* < 0.05; ** *p* < 0.01; *** *p* < 0.001; ns, not significant.

**Extended Data Figure 8. *MPHOSPH8KO* or *MORC2*KO human cerebral organoids show a strong upregulation of repetitive elements and clustered protocadherins.**

**(a)** Immunoblot analysis of two independent clones of control, *MPHOSPH8*KO and *MORC2*KO H9 human embryonic stem cells used for cerebral organoid generation.

The blots were incubated with antibodies against MPP8, MORC2A, LINE-1 ORF1p and β-Actin as loading control.

(**b**) Uniform manifold approximation and projection (UMAP) plot of single-cell RNA-sequencing from day 27 cerebral organoids. Approximately 3400 cells per group were analyzed and are color-coded by genotype. For each group two organoids (each organoid originating from an independent ESC clone) were pooled.

(**c**) Individual UMAP plots for the neuronal precursor marker genes *SOX2*, *PAX6*, *DLX1*, and also *DCX* as marker for immature neurons for all cells shown in (**b**).

(**d**) Representative staining of neural progenitors (SOX2, green) and neurons (HuC/D, red) (upper panels) as well as dorsal neural progenitors (PAX6, green) and the ventral transcription factor (DLX2, red) (lower panels) of day 60 control, *MPHOSPH8*KO and *MORC2*KO cerebral organoids (scale bar = 400 µm). (**e,f,g**) Relative mRNA expression of alpha satellites (**e**), LINE-1 transposable elements (**f**) and *PCDHB2* (**g**) in two independent clones of control, *MPHOSPH8*KO and *MORC2*KO hESCs (day 0) and cerebral organoids (day 25 and day 40). Mean values +/- SD were normalized to the housekeeping gene β-Actin (*ACTB)* (n = 1 for day 0 ESC, n = 3 for day 25 and day 40 cerebral organoids). Data are shown for one of three independent experiments. For panels e-g each data point represents an individual organoid and *p* values were calculated using one-way ANOVA and Dunnett’s multiple comparisons test compared to control A of each time point. ** *p* < 0.01; *** *p* < 0.001; ns, not significant.

## References

1. Kouzarides, T. Chromatin modifications and their function. Cell 128, 693–705 (2007).

2. Black, J. C., Van Rechem, C. & Whetstine, J. R. Histone lysine methylation dynamics: establishment, regulation, and biological impact. Mol Cell 48, 491– 507 (2012).

3. Albert, M. & Helin, K. Histone methyltransferases in cancer. Semin Cell Dev Biol 21, 209–220 (2010).

4. Parkel, S., Lopez-Atalaya, J. P. & Barco, A. Histone H3 lysine methylation in cognition and intellectual disability disorders. Learn Mem 20, 570–579 (2013).

5. Pattaroni, C. & Jacob, C. Histone methylation in the nervous system: functions and dysfunctions. Mol Neurobiol 47, 740–756 (2013).

6. Zhu, Y., Sun, D., Jakovcevski, M. & Jiang, Y. Epigenetic mechanism of SETDB1 in brain: implications for neuropsychiatric disorders. Transl Psychiatry 10, 115–8 (2020).

7. Tchasovnikarova, I. A., et al. GENE SILENCING. Epigenetic silencing by the HUSH complex mediates position-effect variegation in human cells. Science 348, 1481–1485 (2015).

8. Tchasovnikarova, I. A. et al. Hyperactivation of HUSH complex function by Charcot-Marie-Tooth disease mutation in MORC2. Nat. Genet. 49, 1035–1044 (2017).

9. Liu, N. et al. Selective silencing of euchromatic L1s revealed by genome-wide screens for L1 regulators. Nature 553, 228–232 (2018).

10. Albulym, O. M. et al. MORC2 mutations cause axonal Charcot-Marie-Tooth disease with pyramidal signs. Ann Neurol 79, 419–427 (2016).

11. Sevilla, T. et al. Mutations in the MORC2 gene cause axonal Charcot-Marie-Tooth disease. Brain 139, 62–72 (2016).

12. Laššuthová, P. et al. Severe axonal Charcot-Marie-Tooth disease with proximal weakness caused by de novo mutation in the MORC2 gene. Brain 139, e26–e26 (2016).

13. Guillen Sacoto, M. J., et al. De Novo Variants in the ATPase Module of MORC2 Cause a Neurodevelopmental Disorder with Growth Retardation and Variable Craniofacial Dysmorphism. Am J Hum Genet 107, 352–363 (2020).

14. Gu, Z. et al. Silencing of LINE-1 retrotransposons is a selective dependency of myeloid leukemia. Nat. Genet. 53, 672–682 (2021).

15. Lefebvre, J. L., Zhang, Y., Meister, M., Wang, X. & Sanes, J. R. gamma-Protocadherins regulate neuronal survival but are dispensable for circuit formation in retina. Development 135, 4141–4151 (2008).

16. Wang, X. et al. Gamma protocadherins are required for survival of spinal interneurons. Neuron 36, 843–854 (2002).

17. Weiner, J. A., Wang, X., Tapia, J. C. & Sanes, J. R. Gamma protocadherins are required for synaptic development in the spinal cord. Proc. Natl. Acad. Sci. U.S.A. 102, 8–14 (2005).

18. Albertin, C. B. et al. The octopus genome and the evolution of cephalopod neural and morphological novelties. Nature 524, 220–224 (2015).

19. Noonan, J. P., Grimwood, J., Schmutz, J., Dickson, M. & Myers, R. M. Gene conversion and the evolution of protocadherin gene cluster diversity. Genome Res 14, 354–366 (2004).

20. Feng, G. et al. Imaging neuronal subsets in transgenic mice expressing multiple spectral variants of GFP. Neuron 28, 41–51 (2000).

21. Chen, W. V. & Maniatis, T. Clustered protocadherins. Development 140, 3297–3302 (2013).

22. Magklara, A. & Lomvardas, S. Stochastic gene expression in mammals: lessons from olfaction. Trends Cell Biol 23, 449–456 (2013).

23. Toyoda, S. et al. Developmental epigenetic modification regulates stochastic expression of clustered protocadherin genes, generating single neuron diversity. Neuron 82, 94–108 (2014).

24. Lancaster, M. A. et al. Cerebral organoids model human brain development and microcephaly. Nature 501, 373–379 (2013).

25. Lancaster, M. A. & Knoblich, J. A. Generation of cerebral organoids from human pluripotent stem cells. Nat Protoc 9, 2329–2340 (2014).

26. Wang, X., Su, H. & Bradley, A. Molecular mechanisms governing Pcdh-gamma gene expression: evidence for a multiple promoter and cis-alternative splicing model. Genes Dev. 16, 1890–1905 (2002).

27. Wu, Q. et al. Comparative DNA sequence analysis of mouse and human protocadherin gene clusters. Genome Res 11, 389–404 (2001).

28. Lefebvre, J. L., Kostadinov, D., Chen, W. V., Maniatis, T. & Sanes, J. R. Protocadherins mediate dendritic self-avoidance in the mammalian nervous system. Nature 488, 517–521 (2012).

29. Thu, C. A. et al. Single-cell identity generated by combinatorial homophilic interactions between α, β, and γ protocadherins. Cell 158, 1045–1059 (2014).

30. Hajj, El, N., Dittrich, M. & Haaf, T. Epigenetic dysregulation of protocadherins in human disease. Semin Cell Dev Biol 69, 172–182 (2017).

31. Jia, Z. & Wu, Q. Clustered Protocadherins Emerge as Novel Susceptibility Loci for Mental Disorders. Front Neurosci 14, 587819 (2020).

32. Cephalopod Behaviour. (2018).

33. Lewandoski, M., Meyers, E. N. & Martin, G. R. Analysis of Fgf8 gene function in vertebrate development. Cold Spring Harb Symp Quant Biol 62, 159–168 (1997).

34. Tronche, F. et al. Disruption of the glucocorticoid receptor gene in the nervous system results in reduced anxiety. Nat. Genet. 23, 99–103 (1999).

35. Moritz, M. S., Tepp, W. H., Inzalaco, H. N., Johnson, E. A. & Pellett, S. Comparative functional analysis of mice after local injection with botulinum neurotoxin A1, A2, A6, and B1 by catwalk analysis. Toxicon 167, 20–28 (2019).

36. Bankhead, P. et al. QuPath: Open source software for digital pathology image analysis. Sci Rep 7, 16878–7 (2017).

37. Schindelin, J., et al. Fiji: an open-source platform for biological-image analysis. Nat Methods 9, 676–682 (2012).

38. Edgar, R., Domrachev, M. & Lash, A. E. Gene Expression Omnibus: NCBI gene expression and hybridization array data repository. Nucleic Acids Res 30, 207–210 (2002).

39. Hauser, C., Schuettengruber, B., Bartl, S., Lagger, G. & Seiser, C. Activation of the mouse histone deacetylase 1 gene by cooperative histone phosphorylation and acetylation. Mol. Cell. Biol. 22, 7820–7830 (2002).

40. Lancaster, M. A. et al. Guided self-organization and cortical plate formation in human brain organoids. Nat Biotechnol 35, 659–666 (2017).

