## Extended Data Figures 1-8 for "The HUSH complex controls brain architecture and protocadherin fidelity"

### Extended Data Figure 1

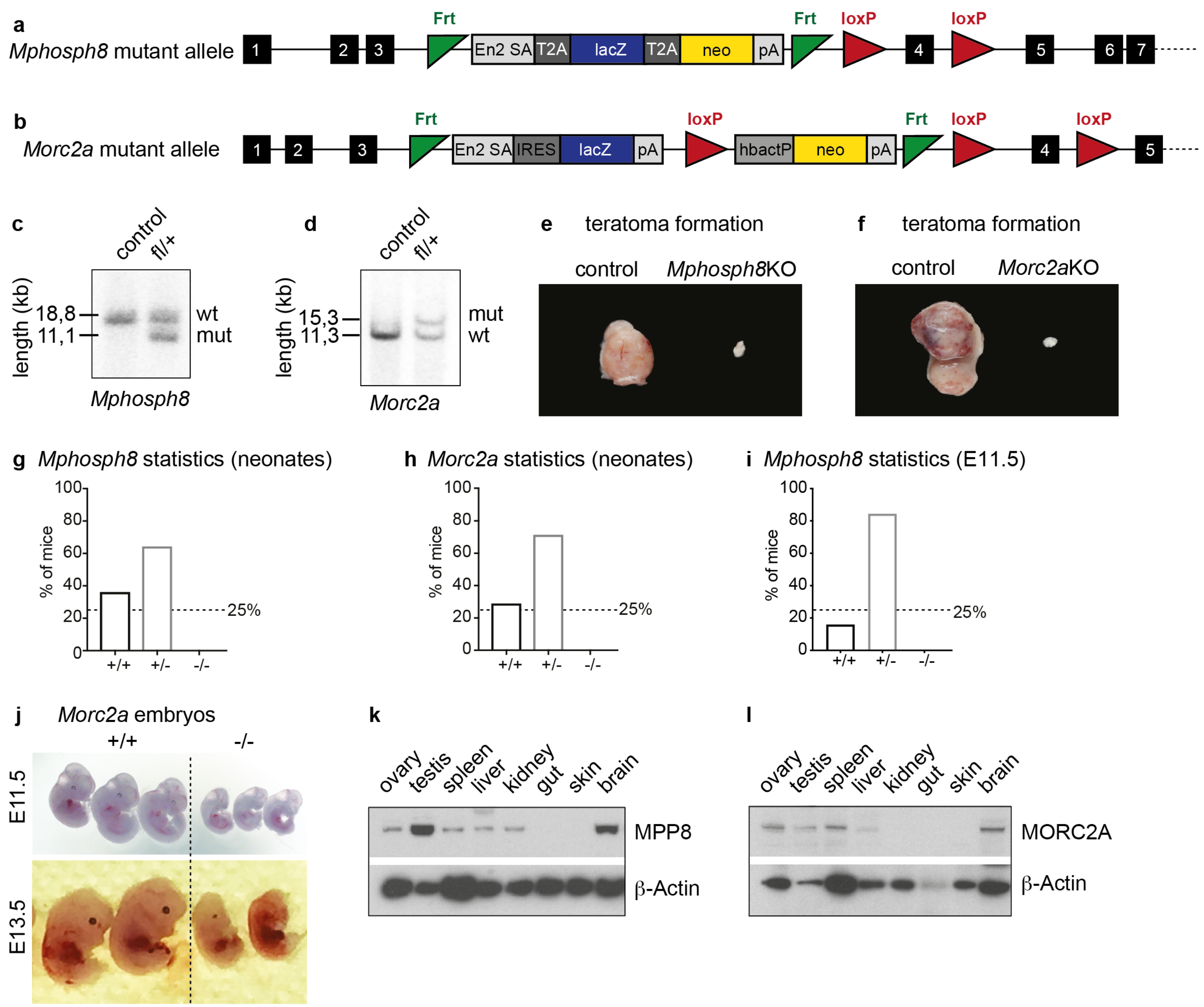

**Extended Data Figure 2**

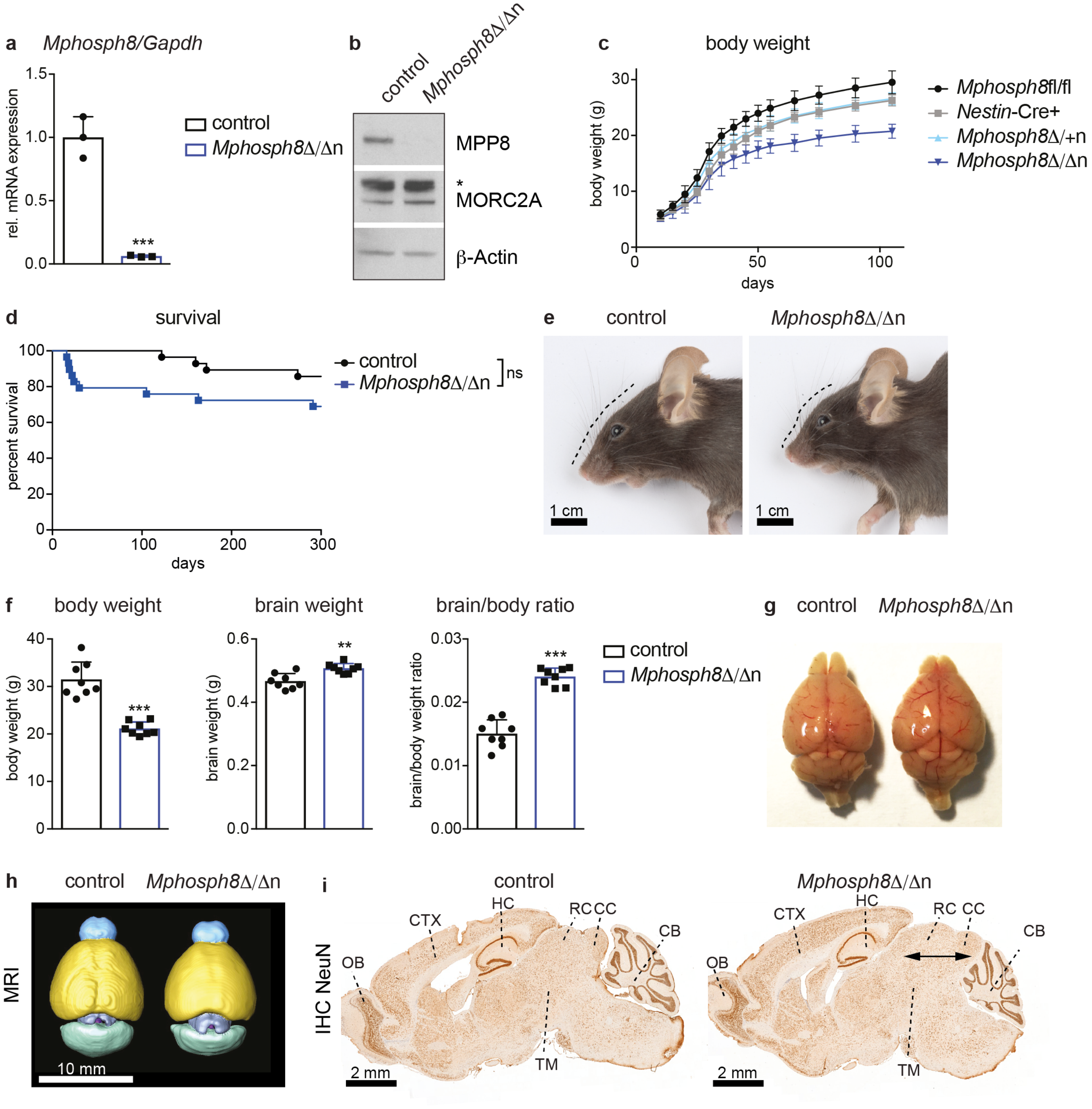

### Extended Data Figure 3

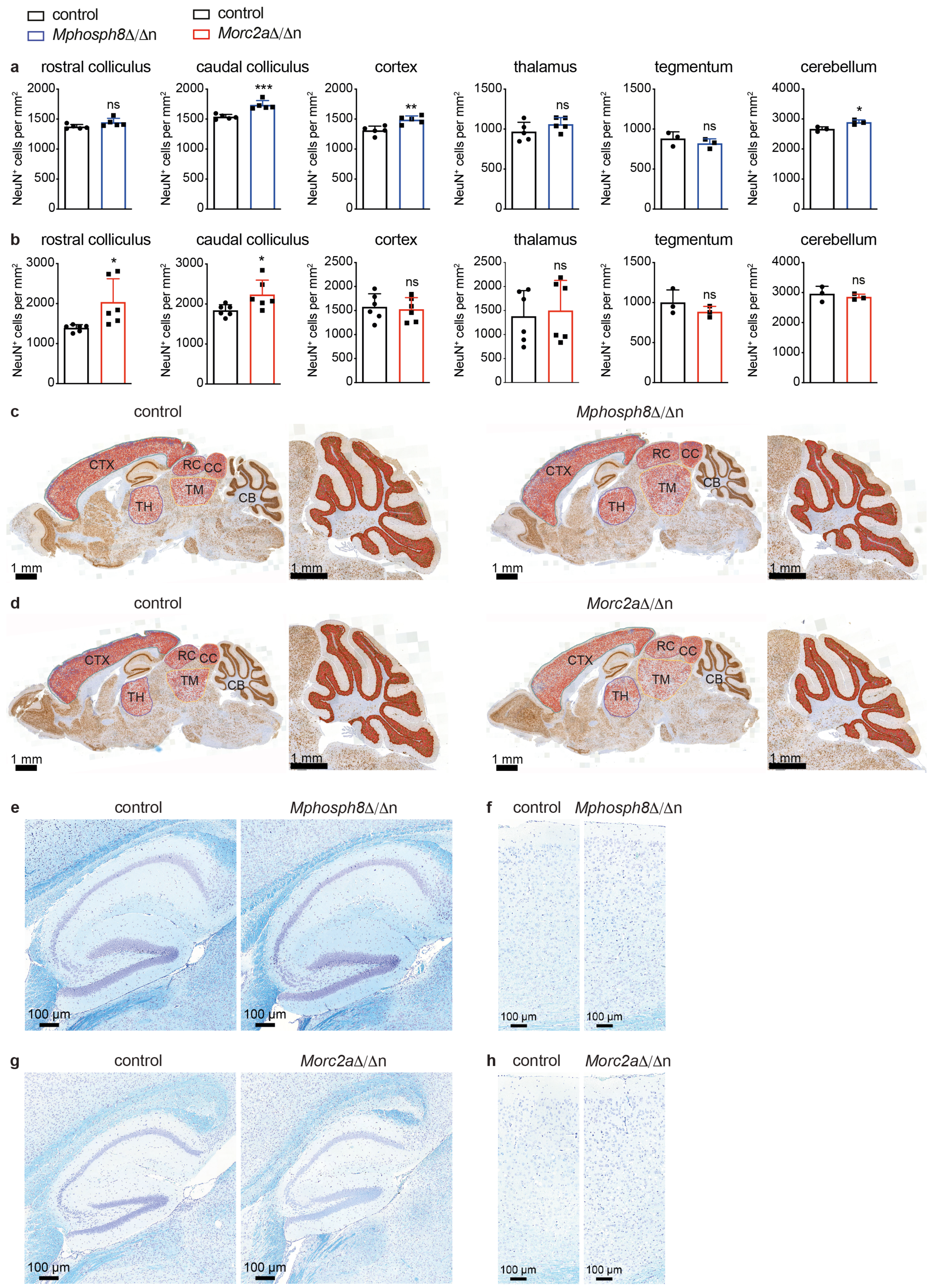

**Extended Data Figure 4**

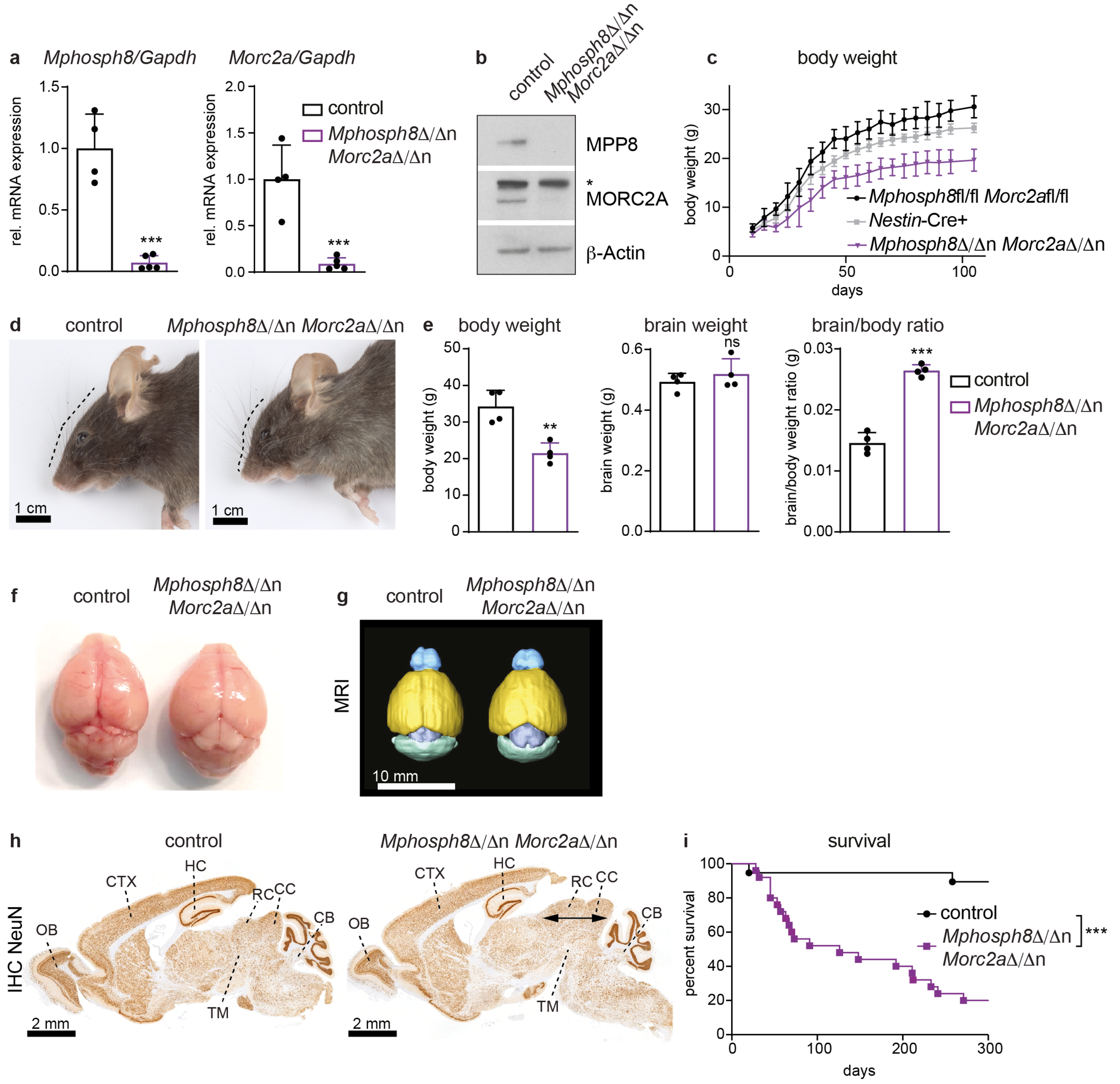

**Extended Data Figure 5**

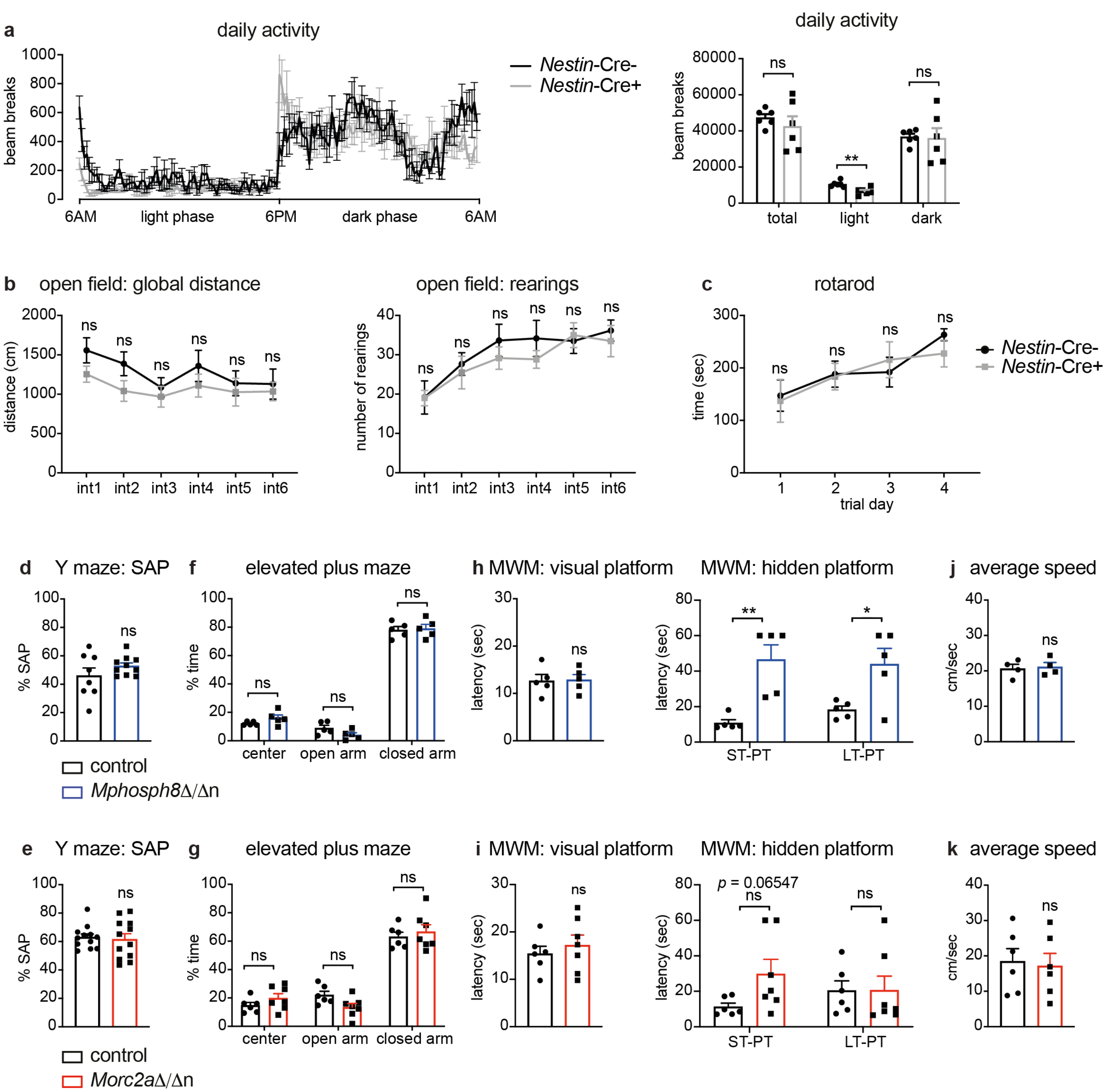

**Extended Data Figure 6**

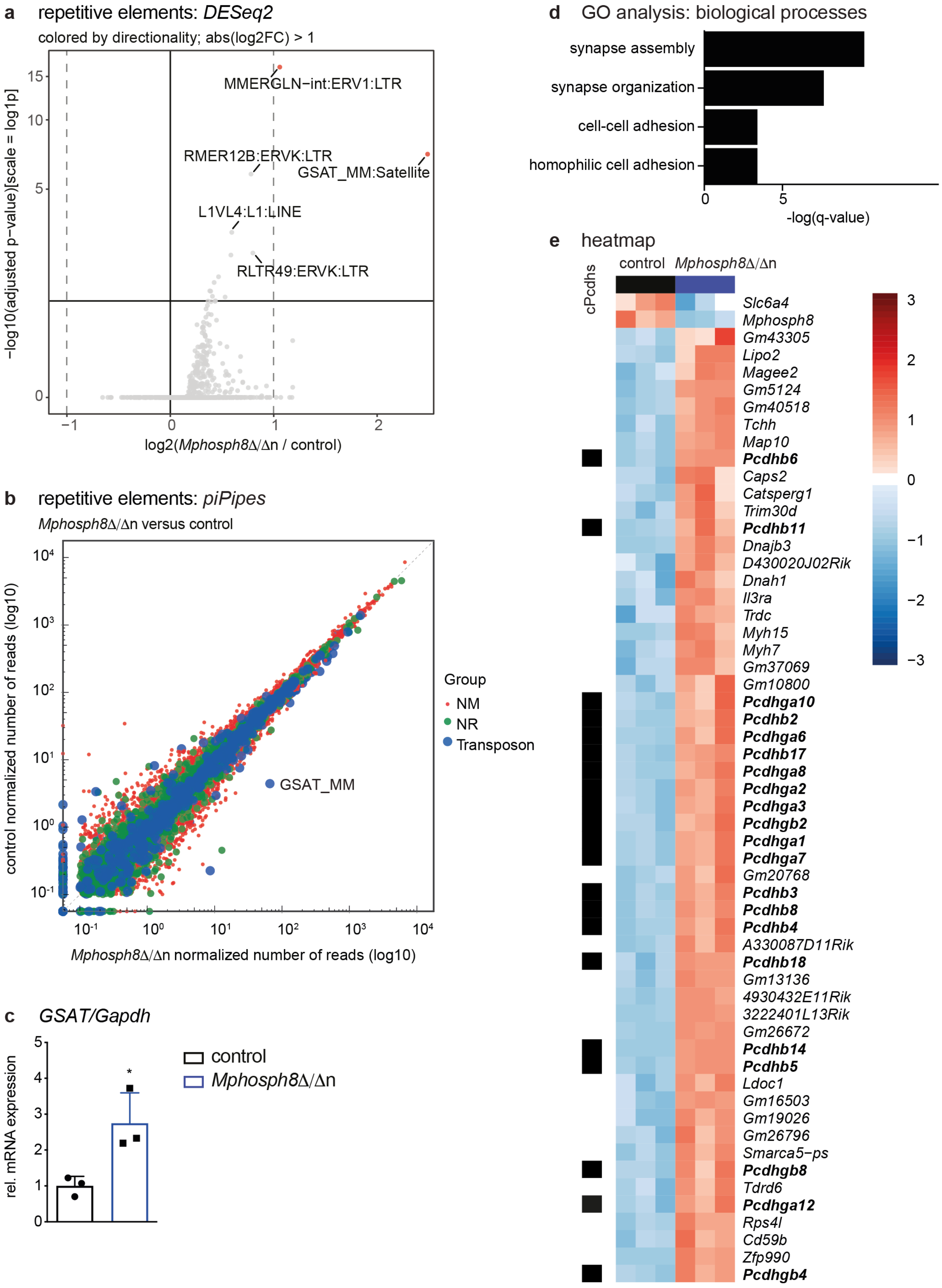

**Extended Data Figure 7**

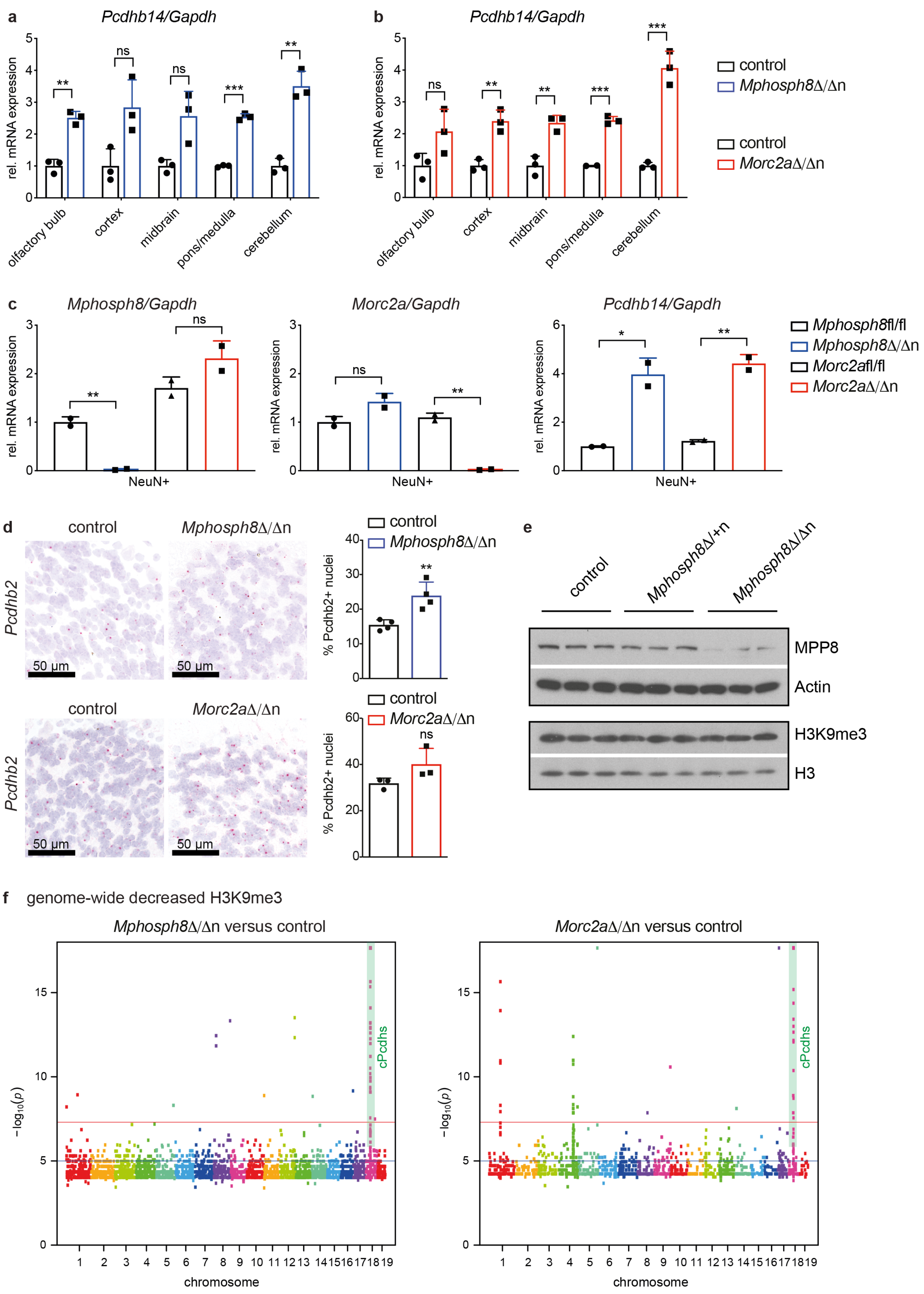

### Extended Data Figure 8

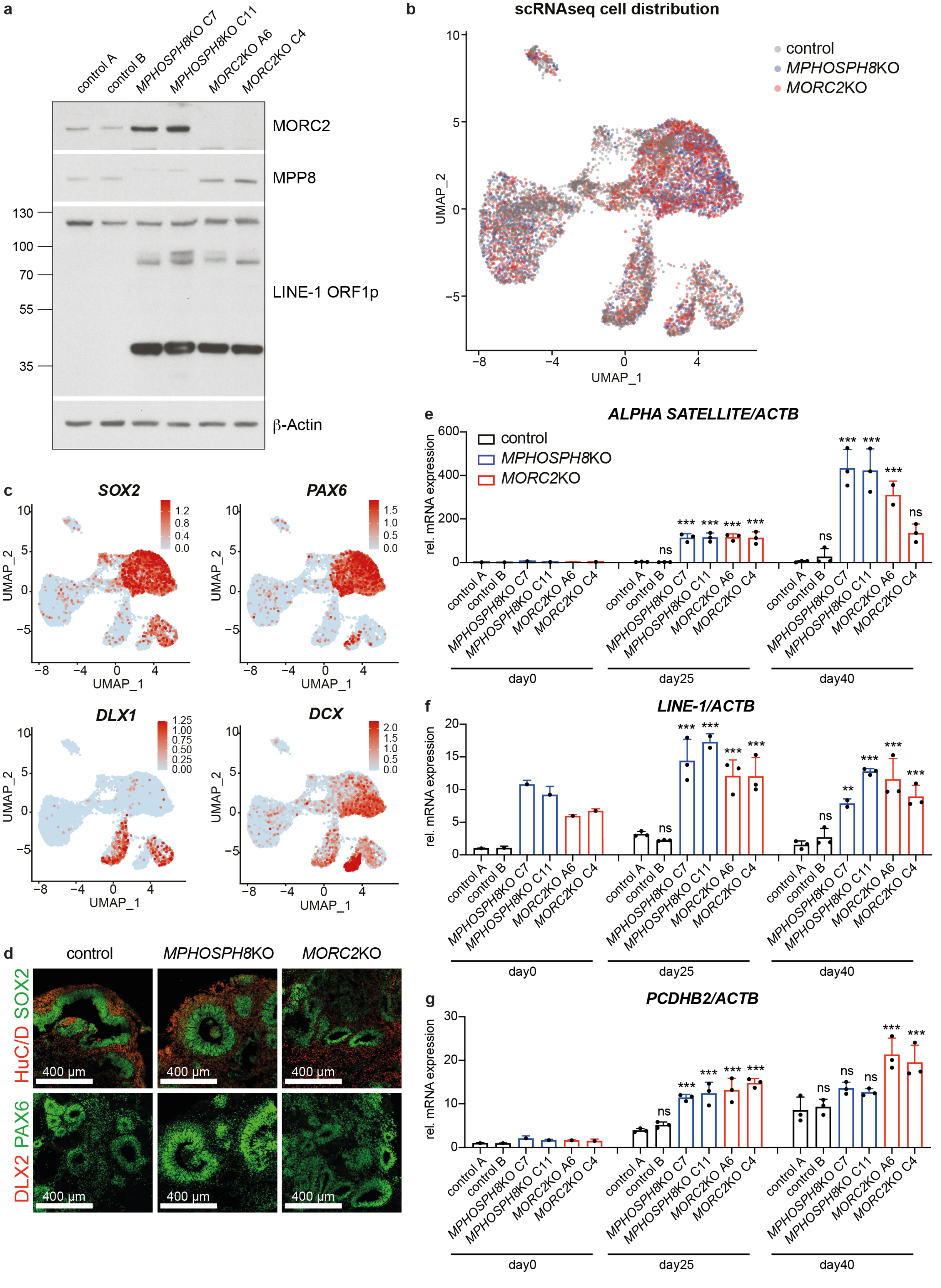
